# Genomic and Kinetic Modeling Involving Nanoparticle-Mediated Delivery of a Novel Chitinase Enzyme to Outpace *Tuta absoluta* Damage

**DOI:** 10.64898/2026.06.23.733920

**Authors:** Yasemin İspirli, Ahmet Can, Mehmet Keçeci, Saliha S. Şahin, Sümeray E. Ayan, Ömür Baysal

## Abstract

The tomato leafminer, *Tuta absoluta*, poses a severe global agricultural threat due to its rapid leaf-mining behavior and swift development of resistance to conventional chemical pesticides. While microbial chitinases are potent biopesticides, their field efficacy is limited by environmental degradation and the short exposure window before larvae penetrate leaf tissues. This study evaluates a stimuli-responsive, controlled-release nanobiopesticide system utilizing a novel chitinase from newly characterized *Serratia marcescens* GBS19. A 61.1 kDa chitinase (GBS19_ChiA) was heterologously expressed in *Escherichia coli* and purified to a specific activity of 215.01 U/mg. The enzyme was immobilized onto starch-coated silica nanoparticles designed for target-triggered release via host alpha-amylase. Genomic profiling and R-based kinetic modeling were integrated to evaluate the efficacy of purified and immobilized forms against *T. absoluta*. Immobilization enhanced thermal and pH stability, with the nanocarrier maintaining 85% activity over 10 weeks. In larval bioassays, immobilization increased mortality from 21.9% to 59.4% (5000 U/mL) by day 3, reaching 62.5% by day 6. Genomic analysis identified an expansive secretome and a Type VI Secretion System (T6SS), characterizing GBS19 as a multi-pronged pathogen. Kinetic modeling established that while immobilized enzymes are effective, the 2.5-hour exposure time on *T. absoluta* requires the synergistic action of chitinases (ChiA/B/C) to reach the lethal desiccation threshold before larvae establish protective mines. Starch-coated silica nanoparticles significantly improve chitinase stability and delivery. However, overcoming the rapid penetration of *T. absoluta* necessitates a whole-cell or multi-enzyme synergistic approach to outpace larval behavioural defences.

## 1. INTRODUCTION

Pesticides play a crucial role in modern agriculture by controlling pathogenic agents and ensuring yield stability. However, only around 0.1% of pesticides applied conventionally reach their intended target, with the remaining 99.9% dispersing into the environment.^1^ The low distribution efficiency of pesticides in the field is primarily due to leaching, runoff, ultraviolet (UV) and rapid evaporation. These factors increase the cost of pesticide application and lead to adverse environmental impacts.^2^

The enzymes produced by microorganisms that are human, animal, or plant pathogens can be highly effective. However, it is possible to utilize these effective enzymes of harmful microorganisms in a beneficial way through recombinant DNA technology. Thanks to cloning technology, the protein produced by the target gene can be artificially expressed, purified, and directed against the target pest.^3^ Currently, there is an increasing focus in research on preparing pesticides as biocompatible, controlled-release systems to enhance their durability and efficiency.^4^ These systems are based on the principle of loading pesticides onto biocompatible materials using immobilisation processes, whereby the pesticide is released over time through diffusion.^5, 6^ This approach prevents losses associated with leaching, runoff, UV degradation and rapid evaporation of pesticides applied in their free form. Within this scope, various nanomaterials that can be utilised for controlled pesticide release have been developed in recent years, employing different active ingredients and enzymes.^7-16^

Among these systems, biodegradable and biocompatible silica nanoparticles have drawn significant attention. This material can also serve as a glutathione-stimulating factor in living organisms.^15, 17, 18^ Glutathione (γ-glutamyl-L-cysteinylglycine), an antioxidant thiol, is widely present in living cells.^19^ In stimuli-responsive pesticide release systems, they have been used to cleave disulfide bonds in drug carriers, thereby releasing the active component.^20-22^ Furthermore, the biodegradability of these silica-based materials arises from the susceptibility of the surrounding disulfide bonds to cleavage in reducing environments.^23^ Owing to their mesoporous structure, these materials do not cause diffusion limitations during drug release or enzyme immobilization.^17^

Considering these factors, silica nanoparticles were chosen as carriers for the immobilization of the chitinase enzyme prepared as a biopesticide in this study. To increase their mechanical stability, these nanoparticles were coated with starch, which is a biodegradable polymer. This choice was made because α-amylase, found in pathogenic organisms, can catalyze the biodegradation of starch by breaking its α-1,4 glycosidic bonds.^24, 25^ Chitinases are highly effective biopesticides as they degrade chitin, a primary structural component of insect cuticles and fungal cell walls, thereby physically disrupting the lifecycle of target pests. The specific chitinase enzyme utilized in this study was previously cloned from *Serratia marcescens* GBS19, heterologously expressed, and biochemically characterized by our research group.^26^ Therefore, the aim of this study is to immobilize the cloned S. marcescens chitinase onto synthesized starch-coated silica nanoparticles and evaluate the formulation’s bioefficacy against significant agricultural pests, specifically the devastating tomato leafminer, *Tuta absoluta* (Meyrick) (Lepidoptera: Gelechiidae).

## 2. MATERIALS AND METHODS

### 2.1. Cloning and Expression of the Chitinase Gene

The chitinase gene from *Serratia marcescens* (GBS19) classified as *Serratia sarumanii* in previous study^27^ was isolated, cloned, and expressed as previously described in detail by.^26^ A 61.1 kDa chitinase (GBS19_ChiA) was amplified from GBS19 genomic DNA using specific primers. Briefly, the isolated gene was inserted into the pET-22b (+) expression vector, and the recombinant plasmid was transformed into competent *Escherichia coli* BL21(DE3) cells. Positive transformants were cultured, and chitinase expression was induced with 0.5 mM IPTG for 4 h at 30°C. The recombinant protein was purified via Ni-NTA affinity chromatography and immobilized onto starch-coated silica nanoparticles to enhance stability for subsequent bioassays.

### 2.2 Precipitation of Ammonium Sulfate by Chitinase Enzyme from Culture Sample

First, both protein and chitinase activity were determined in the supernatant of the culture sample to determine whether the produced chitinase was intracellular or extracellular. Analyses revealed no significant amount of protein ^28^ or chitinase activity in the supernatant. Therefore, it was decided to proceed with the intracellular enzyme purification procedure. For this purpose, the cell pellet obtained from the culture sample was suspended in pH 7.4 (0.1 M) phosphate buffer. Then, sonication at 4°C was applied to lyse the microbial cells. The suspension was separated by centrifugation at 8000 x g for 30 minutes to remove cellular debris. The supernatant containing chitinase was incubated at 40°C for 30 minutes, followed by centrifugation at 12000 x g for 25 minutes to remove insoluble material. Bradford protein determination and chitinase activity assay were performed on the supernatant obtained from this step. The supernatant was partially purified by sequentially precipitation between 25% and 85% ammonium sulfate saturations. At each saturation level, the solution was centrifuged at 12000 x g for 30 minutes, and the resulting protein was subjected to Bradford protein assay and chitinase activity test to determine the optimal ammonium sulfate saturation. After ammonium sulfate precipitation, partial purification was completed by dialyzing the crude protein overnight in a solution containing 50 mM phosphate buffer (pH 7.0) and 10 mM NaCl.^29^

#### 2.2.1 Chitinase activity assay and Purification

0.1 ml of chitinase solution was added to 0.9 ml of 1% (w/v) colloidal chitin, and the mixture was incubated at 55°C for 1 hour. The reaction was stopped by adding 3 ml of DNS (Dinitrosalicylic Acid) and heated at 100°C for 5 minutes. Finally, centrifugation was performed, the reducing sugar DNS method was applied, and absorption measurements were taken at 540 nm using a spectrophotometer (Thermo Scientific-Multiscan Go 510).^30^

The concentrated protein sample was purified by gel filtration on a Sephadex G-100 column (2.5 x 45 cm) pre-equilibrated with 50 mM phosphate buffer (pH 7.0). 5 ml of concentrated protein was loaded into the column, and 5 ml fractions were collected using a peristaltic pump at a flow rate of approximately 30 mL/h. Thirty-four fractions were obtained, and Bradford protein assay and chitinase activity assay were performed on each fraction.^31^

The column (2.5 x 30 cm) was filled with diethylaminoethyl (DEAE) Sephadex A-50 and equilibrated with 0.02 M phosphate buffer (pH 7.0). The fractions with the highest specific activity obtained from gel filtration were loaded onto the DEAE-Sephadex A-50 column. Elution was performed first with 0.05 M phosphate buffer, then with the addition of 0.05 M NaCl in the same buffer at a flow rate of 60 mL/h. The fractions were collected, and Bradford protein assay and chitinase activity assay were performed on each fraction.^31^

The molecular weight of the chitinase enzyme was determined using the SDS-PAGE method described by Laemmli.^32^ Analysis was performed using 10% (w/v) acrylamide in gels. For molecular weight determination, standard protein markers and sample protein bands were stained and visualized with Coomassie Brilliant Blue R-250.^29^ For SDS-PAGE, Thermo Scientific Unstained Protein Molecular Weight Marker, consisting of 6 different proteins between 14.4-116 kDa, was used (β-galactosidase (116 kDa), bovine serum albumin (66.2 kDa), ovalbumin (45 kDa), lactate dehydrogenase (35 kDa), REase Bsp981 (25 kDa), β-lactoglobulin (18.4 kDa) and lysozyme (14.4 kDa).

### 2.3. Synthesis and Characterization of Starch-Coated Silica Nanoparticles

*-OH modification*: 960 mg cetyl-trimethyl-ammonium bromide (CTAB), 31 mL ethanol, 0.5 mL triethylamine, and 188 mL pure water were mixed and incubated at 80°C for 1 hour. 4 mL tetraethyl orthosilicate (TEOS) and 1.6 mL of a solution containing bis(triethoxysilylpropyl) disulfide were added dropwise to this mixture. The mixture was incubated at the same temperature for 6 hours. The precipitate formed at the end of the reaction was centrifuged at 20,000 rpm for 30 minutes and then washed three times with methanol. The resulting precipitate (Precipitate I) was dried.^23, 33^

*-NH_2_ modification*: Precipitate I was suspended in 100 mL of toluene, and the suspension was heated at 90°C for 3 hours with continuous stirring. After the process was complete, the mixture was cooled to room temperature. 750 µl of (3-aminopropyl) trimethoxysilane was added to the suspension, and it was incubated first at room temperature for 1 hour, then at 90°C for 36 hours. At the end of the incubation period, the resulting precipitate was separated by centrifugation and washed three times with methanol. The resulting precipitate (Precipitate II) was dried.

-*COOH modification*: 600 mg of Precipitate II was suspended in 120 mL of acetone using ultrasonic dispersion. Then, 60 mL of 1.5 M succinic anhydride was slowly added to the suspension and mixed. The resulting precipitate was separated by centrifugation and washed three times with ethanol and dried under vacuum (Precipitate III).

*Starch coating*: 500 mg of Precipitate III was homogenously suspended in 50 mL of pH 7.0 Tris-HCl buffer. Then, 400 mg of N-(3-dimethylaminopropyl)-N′-ethylcarbodiimide hydrochloride (EDC) and 300 mg of N-hydroxysuccinimide (NHS) were added to the mixture and stirred for 30 minutes. Subsequently, 60 mL of modified starch solution was added to this mixture, and the mixture was incubated overnight at room temperature with continuous stirring. After the process was complete, the silica nanoparticles coated with modified starch were separated by centrifugation and washed three times with deionized water.

The surface groups and chemical structures of starch-coated silica nanoparticles were investigated using ATR-FTIR analysis (Thermo Scientific Nicolet iS-5 ATR/FTIR Spectrometer). Thermogravimetric analysis (TGA) was performed on a Perkin Elmer TGA 4000 instrument under an N₂ atmosphere, in the temperature range of 50-650°C and at a heating rate of 25°C/min to determine thermal properties. The surface morphology of the nanoparticles was observed after each modification step using a Hitachi Regulus 8230 SEM instrument. Furthermore, particle size distribution and zeta potential measurements were performed after each modification at pH 4.5 and pH 7.0 conditions using a MALVERN Nano ZS90 zeta-sizer.

### 2.4. Optimization and Characterization of Chitinase-Immobilized Starch-Coated Silica Nanoparticles

Purified lyophilized chitinase enzyme was immobilized by adsorption onto starch-coated silica nanoparticles followed by cross-linking with glutaraldehyde. Chitinase amount (0.5, 0.75, 1.0, 1.5, and 2.0 mg/mL), nanoparticle amount (5; 7.5; 10; 12.5 mg/mL), adsorption time (10, 15, 20, and 30 minutes), and glutaraldehyde concentration (1%, 2%, 3%, and 4% (v/v)) were selected as optimization parameters. All experiments were performed in triplicate.

Commercial chitinase enzyme (Chitinase from *Streptomyces griseus*) was obtained from Sigma-Aldrich to compare the enzymatic properties of purified chitinase and immobilized-purified chitinase. All enzymatic properties were studied separately for the three different chitinases. For all parameters in enzymatic properties, the terms free chitinase, immobilized chitinase, and commercial chitinase were used. In this context, optimum temperature (15-60°C), optimum pH (pH 3.0-10.0), thermal stability (15-60°C), pH stability (pH 3.0-10.0), and kinetic parameters (K_m_ and V_max_) were determined. In addition, reusability and storage stability parameters, which are additional features for immobilized chitinase, were also studied.^34^

### 2.5. In vitro applications and calculation of kinetics of a controlled biopesticide release system

Experiments were conducted in distilled water and under optimum conditions, as is applied in insecticide release studies in the literature. For this purpose, chitinase-loaded nanoparticles were placed in a dialysis bag and incubated in 100 mL of 1% chitin solution at 35°C with continuous stirring. At specific intervals, 2 mL samples were taken from the dialysis solution and replaced with the same amount of fresh chitin solution. Chitinase activity was measured in the collected samples to analyze chitin conversion kinetics. Chitin degradation (conversion) kinetics were evaluated using Korsmeyer-Peppas, Higuchi, zero-order and first-order models.^35, 36^

### 2.6 Tomato moth rearing

Mesh cages measuring 50 × 50 × 100 cm (width × length × height) were used for production. Each cage contained four tomato plants grown in pots for approximately six weeks. About 100 adult tomato moths were placed inside the cages and kept for 24–48 hours to allow egg laying. The experiments were conducted in climate chambers set to 25°C, 60% relative humidity, and 16 hours of light.^37-39^

### 2.7 Insecticidal activity trials

For the trials, 2–5-day-old tomato moth adults were collected from the stock culture and allowed to lay eggs on plants for 24 hours, after which the adults were removed. The trials were conducted in 3 cm diameter petri dishes. Agar (1–2%) was poured into the petri dishes, and 3 cm diameter leaf discs obtained from the leaves of insecticide-free plants were placed on the agar surface. Ten eggs less than 24 hours old were then placed in each petri dish using a brush.^40^

In the experiments, biopesticide preparations were made using purified chitinase enzyme or five different doses of chitinase enzyme immobilized on nanoparticles (250, 500, 1000, 2000 and 5000 U/ml) at a concentration of 2.5 mM.

The treatments were conducted using a Potter spray tower (Burkard Manufacturing Co. Ltd.) at a pressure of 0,70 kg/cm². Each Petri dish was individually treated with 1 mL of solution, and the amount applied to each dish was weighed and recorded immediately.^41^ The petri dishes were then placed in climate chambers at 25 ± 1°C, with a relative humidity of 60 ± 10%, and 16 hours of light. At the end of the eighth day, larval hatching from the eggs was checked, and the eggs were classified as live or dead using a stereo microscope.^42^ The experiment included 8 replicates, with each Petri dish considered as one replicate.^40^ The efficacy of purified chitinase enzyme and chitinase-immobilized starch-coated silica nanoparticles against *T. absoluta* larvae was determined using method number 022, as recommended by the IRAC (Insecticide Resistance Action Committee).

In the experiment, leaves from tomato plants grown without insecticide were used. The leaves were cut into discs with a 3 cm diameter, dipped in a chitinase solution, and then left to dry under laboratory conditions. The discs were then placed in petri dishes containing agar, with the lower side of the leaf facing upwards.^37^ Ten second-stage (L2) tomato moth larvae were placed in each petri dish using a brush, and the top of the petri dish was covered with plastic wrap. The plastic wrap was punctured with a needle for ventilation, and the activity of the larvae was monitored.^37^

The experiment was conducted with 32 replicates for each treatment (one larva per replicate). The petri dishes were placed in climate chambers at 25 ± 1°C, 60 ± 10% relative humidity, and 16 hours of light. After 72 hours, the larvae in the petri dishes were recorded as dead or alive. After this period, as the leaf discs on the agar lost their viability, the live larvae were transferred to fresh, untreated leaf discs. A second count was performed at the end of the sixth day. Larvae that did not react when touched with a brush were considered dead.^38^

In the experiments, distilled water was used as a negative control and emamectin benzoate (5.7% WG) insecticide at a dose of 40 g/100 L was used as a positive control. During the preparation of the solutions, g/L Triton X-100 was added to the distilled water.

### 2. 8 Bioinformatics analysis

The genome of *Serratia marcescens* GBS19 was annotated using Prokka v1.14.6 to identify protein-coding sequences (CDS). Functional categorization was performed via EggNOG-mapper and the KEGG database. Pathogenicity-associated islands were identified through manual curation and local BLASTp searches against the Virulence Factor Database (VFDB).

To identify extracellular secretations, the GBS19 proteome was screened for signal peptides using SignalP 6.0. Proteins containing a Sec/SPI cleavage site with a probability >0.90 were classified as secreted. The domain architecture of top chitinase candidates (ChiA, ChiB, ChiC) was mapped using InterProScan and the Pfam database to identify catalytic Glycosyl Hydrolase 18 (GH18) domains and Chitin-Binding Modules (CBMs). To compare the efficiency of the treatments, a Kinetic Race Model was developed in R v4.3.0. The model simulated the rate of cuticular erosion (y) as a function of time (t), comparing the synergistic output of the GBS19 triad against the static rate of the cloned enzyme. The success window was defined as the duration required to reach the lethal threshold of desiccation before the documented mean penetration time (2.5 hours). To determine the extracellular localization of identified virulence factors, the GBS19 proteome was also analyzed using SignalP 6.0. Sequences were screened for the presence of Sec/SPI (signal peptidase I) cleavage sites. Proteins were classified as secreted if a signal peptide (SP) was predicted with a probability >0.90. Integration of BLASTp results and SignalP predictions was performed in R v4.3.0 using the dplyr and ggplot2 packages. Biosynthetic Gene Clusters (BGCs) were identified using antiSMASH 7.0 (Bacterial version) with detection rules to capture both well-characterized and novel clusters. Specialized metabolism, including Non-Ribosomal Peptide Synthetases (NRPS) and Polyketide Synthases (PKS), was prioritized based on their known roles in insect immune suppression.

## 3. Data Visualization and Statistical Analysis

All genomic visualizations, including multi-panel violin plots and stacked secretome charts, were generated in R (V.4.4.3). To establish a baseline for comparative homology, reference amino acid sequences for known entomopathogenic virulence factors were retrieved from the NCBI Protein database. Sequence retrieval was automated via the Entrez Programming Utilities (E-utilities) using the rentrez package in R. Search queries were specifically formulated to target well-characterized insecticidal protein families from established entomopathogens, including *Bacillus*, *Serratia*, and *Pseudomonas*, and *Photorhabdus* species. The targeted functional categories included Chitinases (gut barrier degradation), Metalloproteases (host immune suppression), Lipases/Phospholipases (cell membrane lysis), and Cry Toxins (pore formation). The complete predicted proteome of the bacterial strain (provided as a faa FASTA file) was formatted into a local sequence database using the makeblastdb command-line tool. High-confidence identification of virulence factors was performed using the BLASTp algorithm (BLAST+ suite). To ensure strict orthology, initial alignments were filtered using a stringent threshold, retaining only hits with an expected value (E-value) ≤ 1 X 10^-20^ and an amino acid identity > 30%. To prevent the duplicate counting of protein isoforms, the results were aggregated by the subject gene ID, and only the highest-scoring alignment (maximum bit-score) was retained for each candidate. To identify putative distant homologs and potentially novel virulence factors that deviate from standard databases, a low-stringency deep homology scan was executed. The BLASTp parameters were relaxed to an E-value threshold of 1.0 to capture distant evolutionary relatives. The output was subsequently filtered to isolate sequence alignments exhibiting between 15% and 45% amino acid identity. Candidates falling within this homology were classified as putative novel variants and mapped against their predicted entomopathogenic properties.

A theoretical, bipartite interaction network was constructed to model the predicted molecular interface between the bacterial virulence factors and the physiological targets of the insect host (*T. absoluta*). The network was built using the igraph package in R (4.4.3). Directed edges were manually curated based on established Lepidopteran pathology models, linking bacterial effectors (e.g., Cry toxins, ChiA, InhA, PlcA) to their corresponding host receptors or substrates (e.g., Cadherin/ABCC2 receptors, Chitin Synthase 1, Cecropin antimicrobial peptides, and apoptosis-related Caspases).

All data processing, statistical summarization, and graphical rendering were performed within the R statistical computing environment (v4.4.3). Data wrangling and categorical grouping were executed utilizing the dplyr package. Visualizations were generated using ggplot2. To assess the conservation of the bacterial functional genomic property, homology distributions across functional categories were analysed using violin plots overlaid with median boxplots. For complex density visualizations, such as the deep homology maps, non-overlapping textual annotations were applied using the ggrepel package. Colorblind-safe, high-contrast palettes from the viridis and RColorBrewer libraries were utilized across all figures to ensure data clarity.

In the egg toxicity trial, eggs from which larvae hatched were considered alive, while those in which hatching did not occur were considered dead. The mortality rate was calculated from the live and dead counts, and the efficacy rate was determined by applying Abbott’s formula ^43^ to this mortality rate. The difference between purified and immobilized chitinase doses was assessed using analysis of variance, and differences between doses were identified using the Tukey test (p<0.05).

In the larval toxicity trials, the difference between purified and immobilized chitinase doses on days 3 and 6 was determined using non-parametric analysis of variance (Kruskal-Wallis test), and statistical differences were identified using the Tukey test (p<0.05).

## 3. RESULTS AND DISCUSSION

### 3.1. Verification of Chitinase Gene Cloning, Expression and Purification

Successful cloning and expression of the chitinase gene from *S. marcescens* GBS19 in the *E. coli* BL21(DE3) system was confirmed, consistent with previous findings.^26^ The recombinant expression vector was verified by restriction enzyme digestion and next-generation sequencing (NGS), which revealed 100% sequence identity with the target chitinase gene.

A total of 13 fractions were obtained via ammonium sulfate precipitation. Among the fractions obtained at different ammonium sulfate saturations, the highest chitinase activity (3660.55 U) and specific activity (78.46 U/mg protein) were obtained at 65% saturation, while significant chitinase activity (2438.84 U) and specific activity (63.22 U/mg protein) values were also reached at 60% saturation. Therefore, for the next purification step, these two fractions (pellets) were combined and used for gel filtration chromatography. Following ammonium sulfate precipitation, 34 fractions were obtained from a Sephadex G-100 column, and specific activity and total protein analyses were performed. Gel filtration chromatography revealed a single major peak (fraction 16) exhibiting chitinase activity, with a specific activity of 143.92 U/mg protein. The fraction (16th) obtained by gel filtration on a Sephadex G-100 column and showing chitinase activity was chromatographed on a DEAE-Sephadex A-50 anion exchange column, and 32 fractions were obtained. The specific activity and protein analyses of these fractions were performed. Ion exchange chromatography yielded a main peak showing chitinase activity, with a specific activity of 215.01 U/mg protein. All the results obtained in the purification steps are presented in Table 1.

**Table 1.**
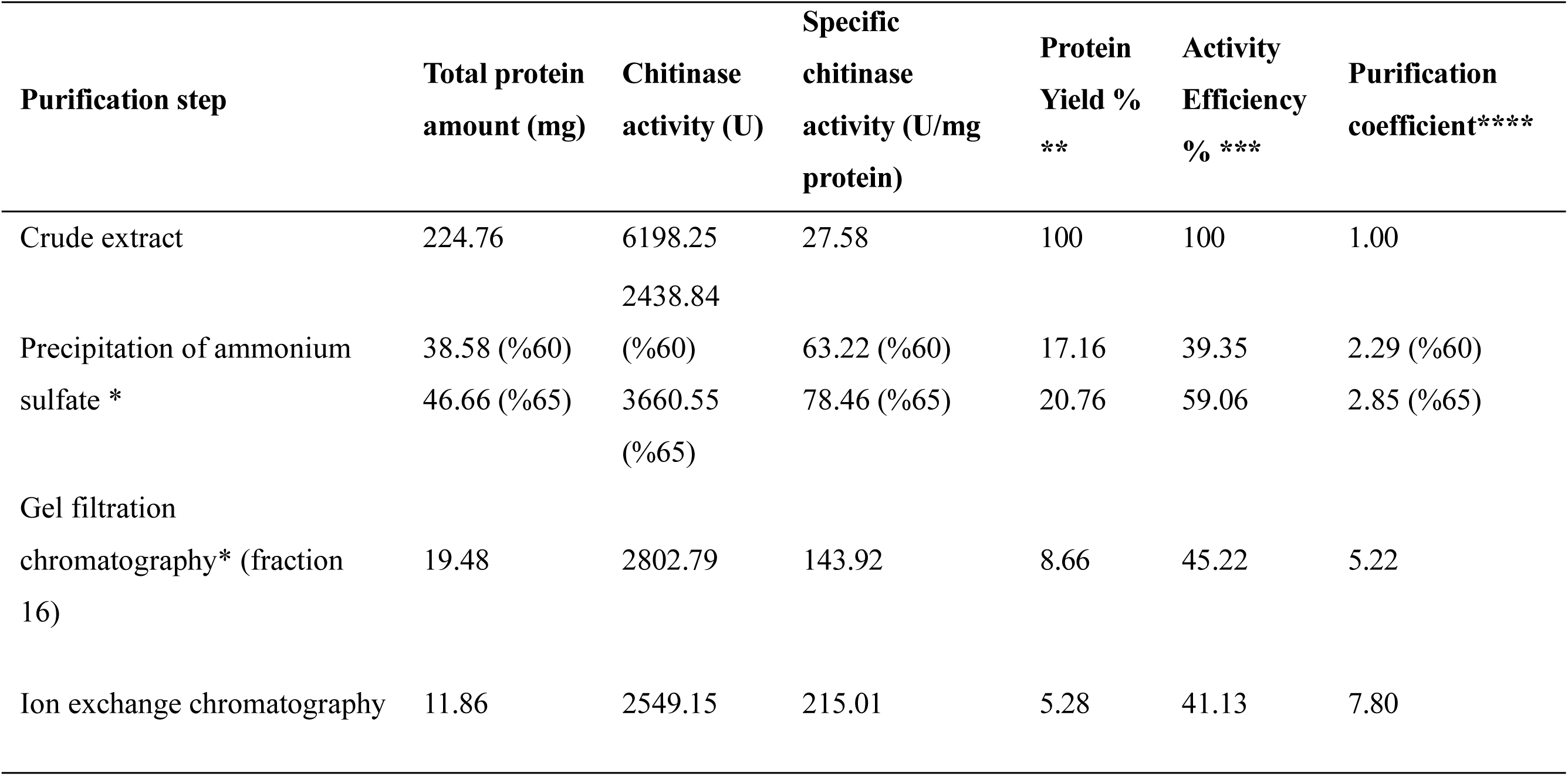
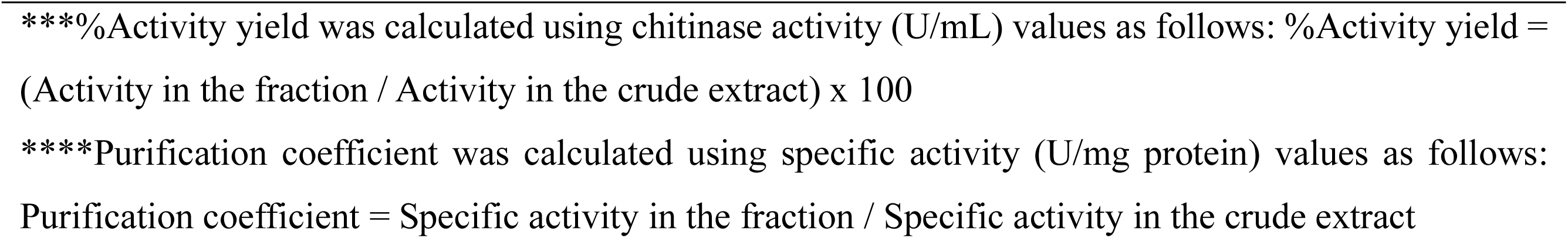
Summary of chitinase purification steps. Table indicates the specific saturation percentages (60% and 65%) used during ammonium sulfate precipitation, and the specific fraction (**Fraction 16**) collected during gel filtration chromatography that were analyzed or used for subsequent steps. Protein yield (%) represents the percentage of total protein recovered at a given step relative to the initial crude extract (set at 100%). *** Activity Efficiency (%) (also known as recovery) represents the percentage of total chitinase activity retained at a given step compared to the initial crude extract (set at 100%). **** Purification coefficient (or purification fold) indicates the relative increase in enzyme purity, calculated by dividing the specific chitinase activity of the current step by the specific chitinase activity of the initial crude extract.

In our study, the chitinase gene region found in *Serratia* genome was transferred to *Escherichia coli*. The data obtained for the purified chitinase were evaluated comparatively with similar studies in the literature. The total activity value of 2549.15 U (specific activity 215.01 U/mg protein) obtained as a result of our purification strategy, which included ammonium sulfate precipitation, gel filtration, and ion exchange chromatography steps, is significantly higher than the 96 U (specific activity 2.34 U/mg protein) value reported in the study with *Stenotrophomonas rhizophila* and the 333.8 U (specific activity 30.3 U/mg protein) value obtained with the transfer of *Paenibacillus barengoltzii* gene region.^44, 45^

The crude extract of chitinase and all purification steps (ammonium sulfate (65%) precipitation, fraction 16 after gel filtration chromatography, and fraction 13 after ion exchange chromatography) were verified using SDS-PAGE. SDS-PAGE band images are given in **Figure 1 (a).** The single band obtained after ion exchange chromatography confirmed that the chitinase enzyme was obtained in pure form. Using the calibration profile based on protein standards given in **Figure 1 (b)**, the molecular weight of the purified chitinase was calculated. As a result of this calculation, the R_f_ value of chitinase was found to be 0.274. Substituting this value into the calibration equation, the molecular weight (M_w_) of the purified chitinase was calculated as 61.1 kDa.

**Figure 1.**
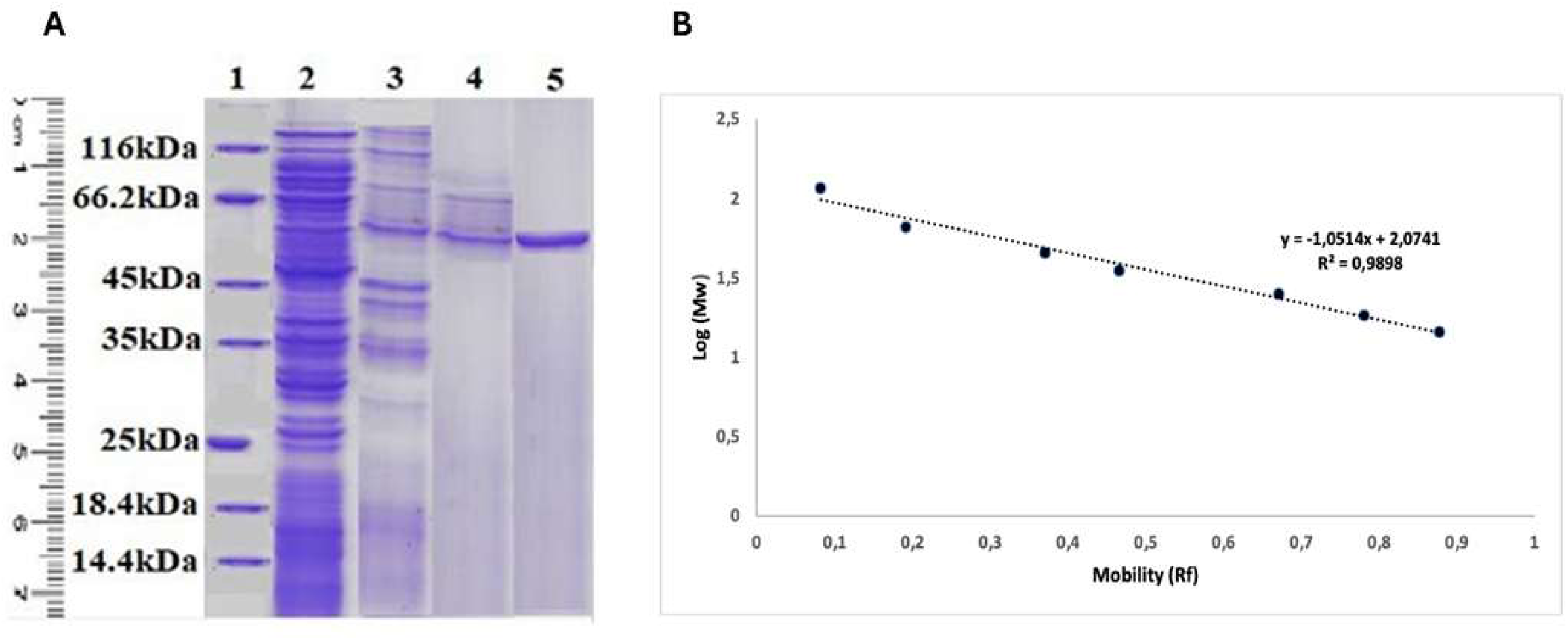
(a) SDS-PAGE band images of chitinase purification steps: 1-Protein marker; 2-Crude extract; 3-Ammonium sulfate (65%) precipitation result; 4-After gel filtration chromatography (fraction 16); 5-After ion exchange chromatography (fraction 13); (b) Calibration profile of standard proteins.

When the M_w_ of the chitinase purified in this study was compared with the molecular weight results of chitinases isolated from microorganisms via direct or recombinant gene transfer in the literature, it was observed that the molecular weight of chitinases purified in the literature is generally in the range of 50-70 kDa. Therefore, the molecular weight calculated as 61.1 kDa in this study was found to be consistent with the literature.^44-49^

### 3.2 Synthesis and Characterization of Starch-Coated Silica Nanoparticles

Surface of silica nanoparticles were successively modified with -OH, -NH_2_, and -COOH, and finally coated with starch. **Figure 2** illustrates the FTIR spectra of silica nanoparticles following modification and coating processes. The spectrum in **Figure 2(a)** corresponds to silica nanoparticles obtained after - OH modification. In this spectrum, the absorption peak observed between 1020–1110 cm⁻¹ is attributed to the Si-O-Si asymmetric stretching vibration, while the peak at 960 cm⁻¹ reflects the asymmetric bending and stretching vibrations of Si-OH. **Figure 2(b)** presents the FTIR spectrum of the -OH modified silica nanoparticles after subsequent -NH₂ modification. The presence of asymmetric deformation vibrations at 1425 and 900 cm⁻¹, characteristic of the amino group, indicates the successful integration of amino groups onto the surface of the silica nanoparticles. **Figure 2(c)** displays the spectrum of the silica nanoparticles subjected to -COOH modification following the -NH₂ and -OH treatments. The vibration peaks at 1725 and 1461 cm⁻¹ demonstrate that the carboxyl group has been integrated into the structure. These findings confirm the successful sequential modification of the silica nanoparticle surfaces. Finally, **Figure 2(d)** shows the FTIR spectrum of the modified silica nanoparticles after starch coating. The observation of primary characteristic bands related to -OH, -CH₂, -CH, and - CO bonds signify the successful formation of the composite structure. The results obtained from the FTIR analysis are in good agreement with similar studies in the literature. According to a study by Socrates,^50^ the peak at 1480 cm⁻¹ is associated with the alkyl group. Similarly, several studies have attributed the peaks at 1040 cm⁻¹, 1220 cm⁻¹, and 960 cm⁻¹ to the asymmetric stretching vibrations of Si–O–Si and Si–O–H bonds.^51-53^ In another study, amino-functionalized silica exhibited the same silica network pattern between 900 and 1200 cm⁻¹; furthermore, the peak at 1560 cm⁻¹ was originated from the -NH₂ group.^54^

**Figure 2.**
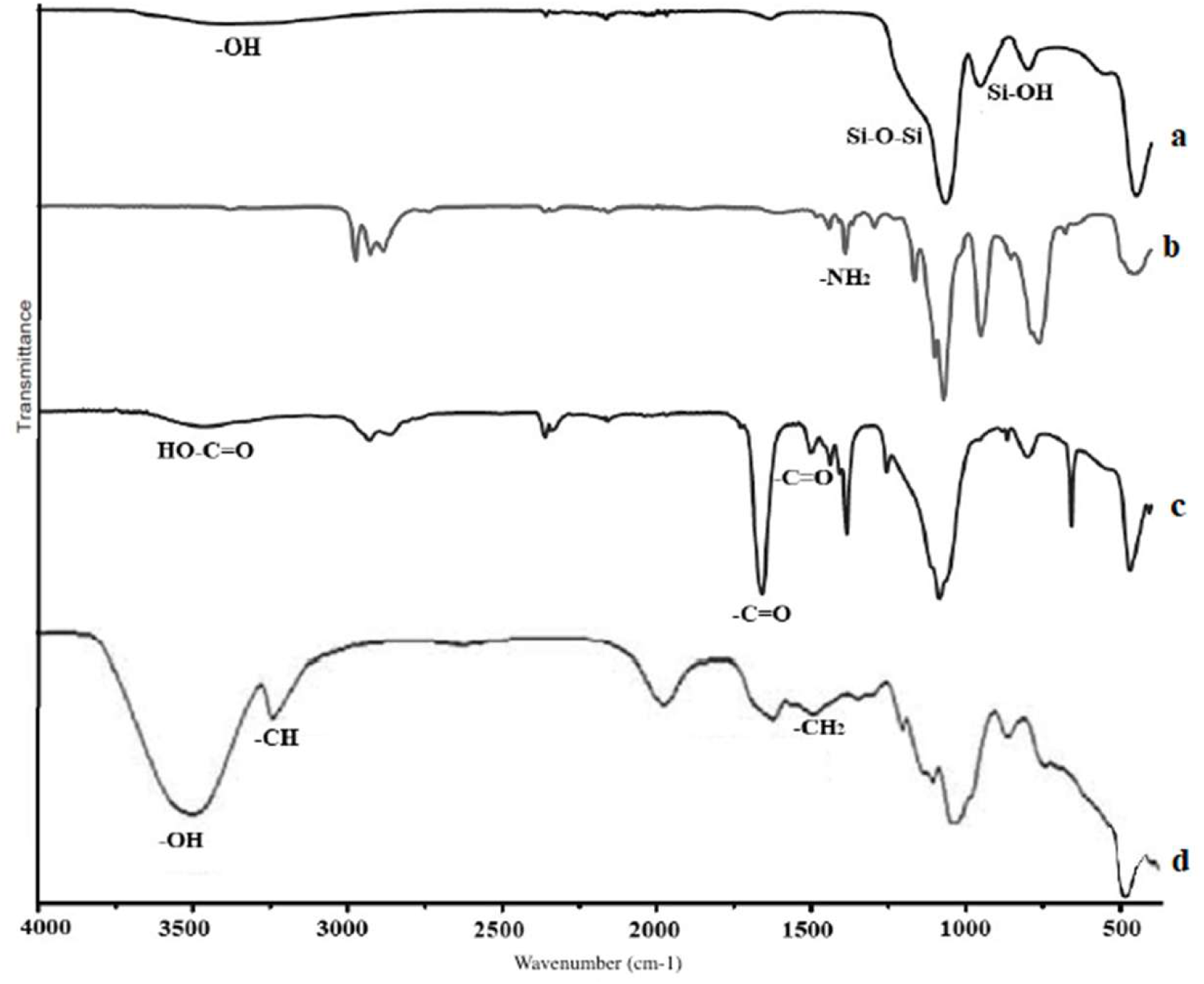
FTIR spectra (a) after -OH modification (b) after -NH_2_ modification (c) after -COOH modification (d) after starch coating

The results of the thermogravimetric analysis (TGA), conducted to verify the starch coating content of the modified structure, are presented in **Figure 3(a)**. Upon examining the TGA thermogram, the initial weight loss occurred at 100°C, which is attributed to the evaporation of structural water and low-molecular-weight compounds.^55, 56^ The second degradation stage took place at approximately 290–300°C and can be assigned to the evaporation of molecular water within the polymeric structure (starch) and the subsequent thermal decomposition of the starch.^56^ Continuous heating led to an increased rate of degradation by weakening the strong intermolecular interactions of the starch. This stage corresponds to the elimination of hydroxyl groups, thermal decomposition, and the depolymerization of the starch carbon chains. The temperatures observed for this step are consistent with those reported for nano-SiO₂-starch films.^57^

**Figure 3.**
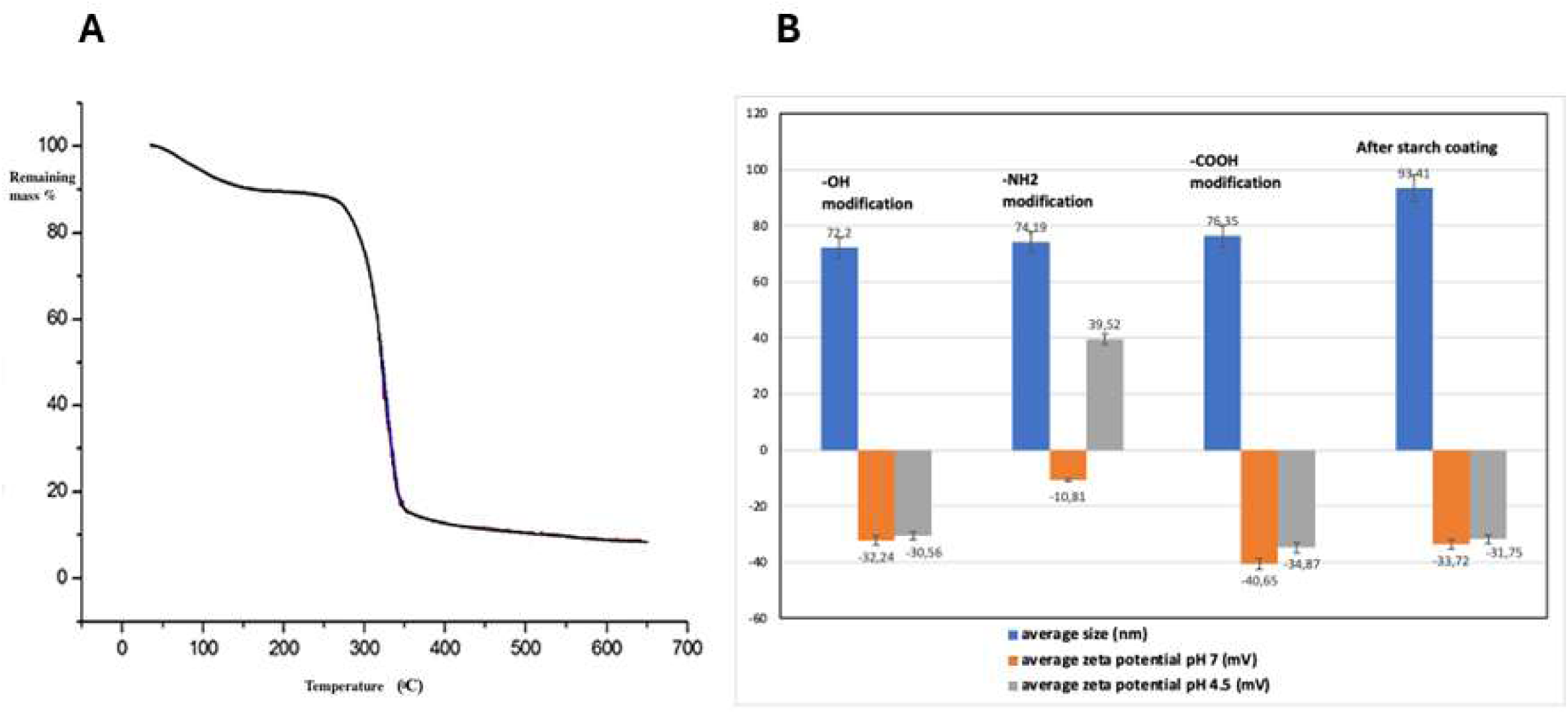
Physical characterization of synthesized nanocarriers. (a) Thermogravimetric analysis (TGA) thermogram tracking the quantitative thermal degradation and mass loss % profile of starch-coated modified silica nanoparticles under an N₂ atmosphere. (b) Combined dynamic light scattering (DLS) hydrodynamic diameter measurements (nm) and surface charge zeta potential alterations (mV) monitored under pH 4.5 and pH 7.0 analytical conditions across sequential functionalization steps.

The average sizes and zeta potentials of the nanoparticles following each modification step are presented in **Figure 3(b)**. No significant change was observed in the average sizes of the nanoparticles after the surface modifications (-OH, -NH₂, and -COOH). This is attributed to the fact that the attached functional groups are not large enough to cause a measurable increase in size. However, a size increase of approximately 20 nm was observed after coating with the starch polymer, as starch is a significantly larger molecule compared to the groups attached during surface modification. This result is consistent with the diameter measurements obtained from the SEM images shown in **Figure 4**. Zeta potential measurements after -NH₂ modification at pH 4.5 clearly demonstrate the surface changes resulting from the reaction with 3-(aminopropyl) trimethoxysilane. Due to the acidic nature of the silanol groups, the silica nanoparticles were negatively charged (-30.56 mV) following -OH modification; however, this value increased to 39.52 mV after the reaction with 3-(aminopropyl)trimethoxysilane. At pH 4.5, primary amines undergo protonation, leading to a high positive zeta potential. Conversely, at pH 7.0, the zeta potential was measured as -10.81 mV, as amines are only partially protonated at neutral pH.

**Figure 4.**
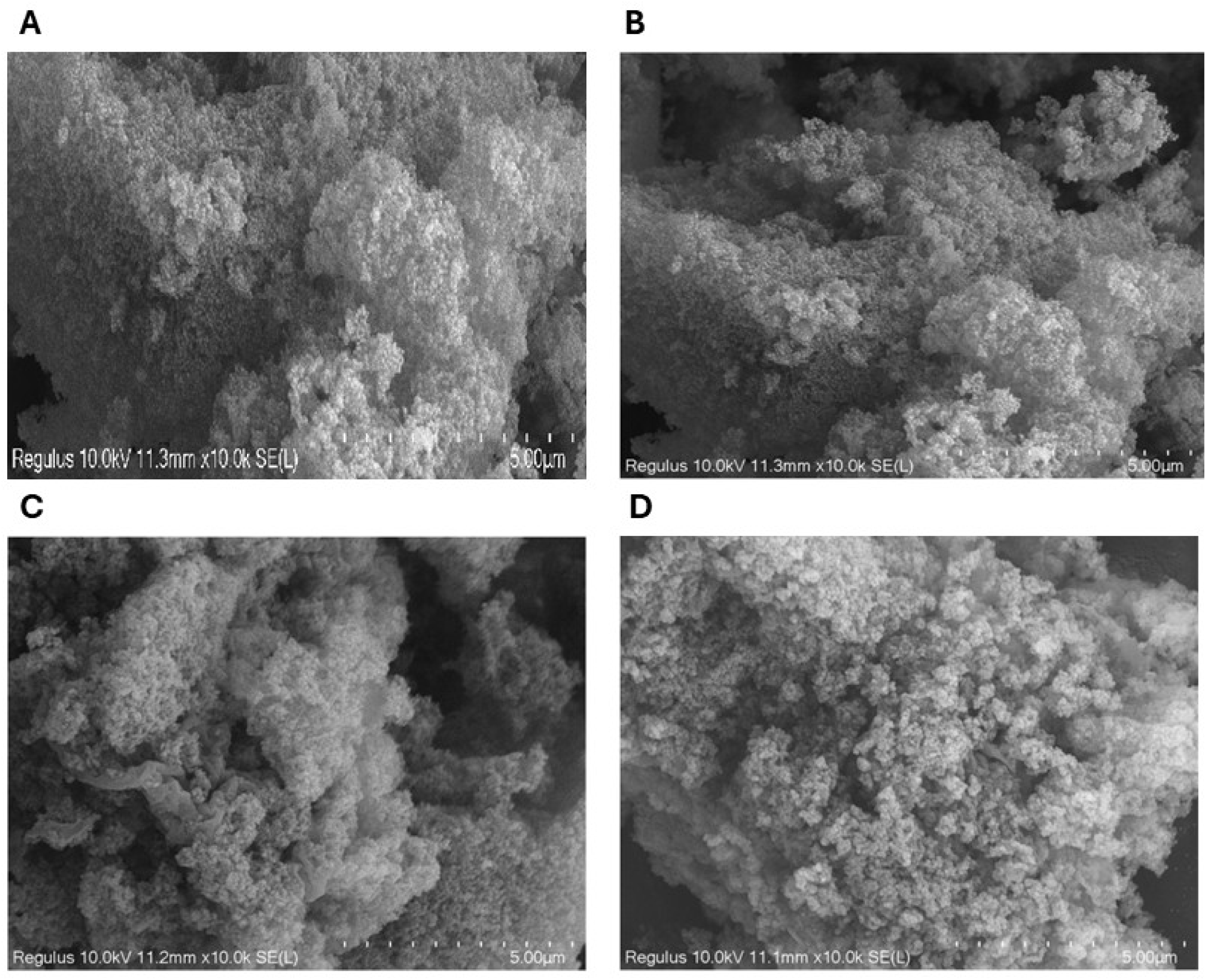
Scanning Electron Microscopy (SEM) morphological micro-topography. High-resolution structural morphology views of synthesized silica nanoparticles tracking physical matrix integrity across processing steps: (a) Highly uniform monodisperse pristine silica core spheres following baseline -OH treatment; (b) Well-dispersed structural matrix following amino (-NH₂) functionalization; (c) Intact structural parameters following carboxyl (-COOH) succinic anhydride derivatization; (d) Final composite assemblies following target polysaccharide structural starch encapsulation showing minor surface matrix bridging. Magnification = 10.0k times; context scale lines indicate 5.00 µm across all panels.

The morphological characteristics of the silica nanoparticles following modifications were examined using scanning electron microscopy (SEM), and the SEM images are presented in **Figure 4**. These images reveal that the modified silica nanoparticles possess a spherical shape and are approximately uniform in size. In **Figure 4(a), 4(b),** and **4(c)**, no significant aggregation was observed among the nanoparticles after -OH, -NH₂, and -COOH surface modifications, respectively. However, after starch coating (**Figure 4 (d)**), it was observed that the nanoparticles began to form aggregates due to the influence of the starch; nevertheless, this aggregation was not extensive and did not significantly increase the nanoparticle size. A rough surface and porous structure were visible in all SEM images, indicating that the modified silica nanoparticles provide an ideal surface for immobilization. Following surface modifications, no remarkable change was observed in the average diameters of the nanoparticles, which remained at approximately 75 nm. After starch coating, an increase in average diameter occurred due to the aggregation, with the average diameter measured at 90 nm. These surface characteristics and average diameter values are consistent with SEM studies on silica nanoparticles reported in the literature.^58, 59^

### 3.3. Optimization and Characterization of Chitinase-Immobilized Starch-Coated Silica Nanoparticles

For the immobilization of chitinase onto starch-coated silica nanoparticles, the chitinase concentration was initially optimized, and the results are presented in **Table 2**. The optimum chitinase concentration was determined to be 1.5 mg/mL, while keeping the nanoparticle concentration at 10 mg/mL, the adsorption time at 20 minutes, and the glutaraldehyde concentration at 2% constant. Although there are limited studies regarding chitinase immobilization in the literature, a few examples are available. In one of these studies, chitinase partially purified from *Trichoderma harzianum* CECT2413 was immobilized onto magnetic nanoparticles. In that study, the optimum chitinase amount was determined to be 6.2 mg/g carrier, resulting in an activity value of 21.48 U at this enzyme loading.^60^

**Table 2.**
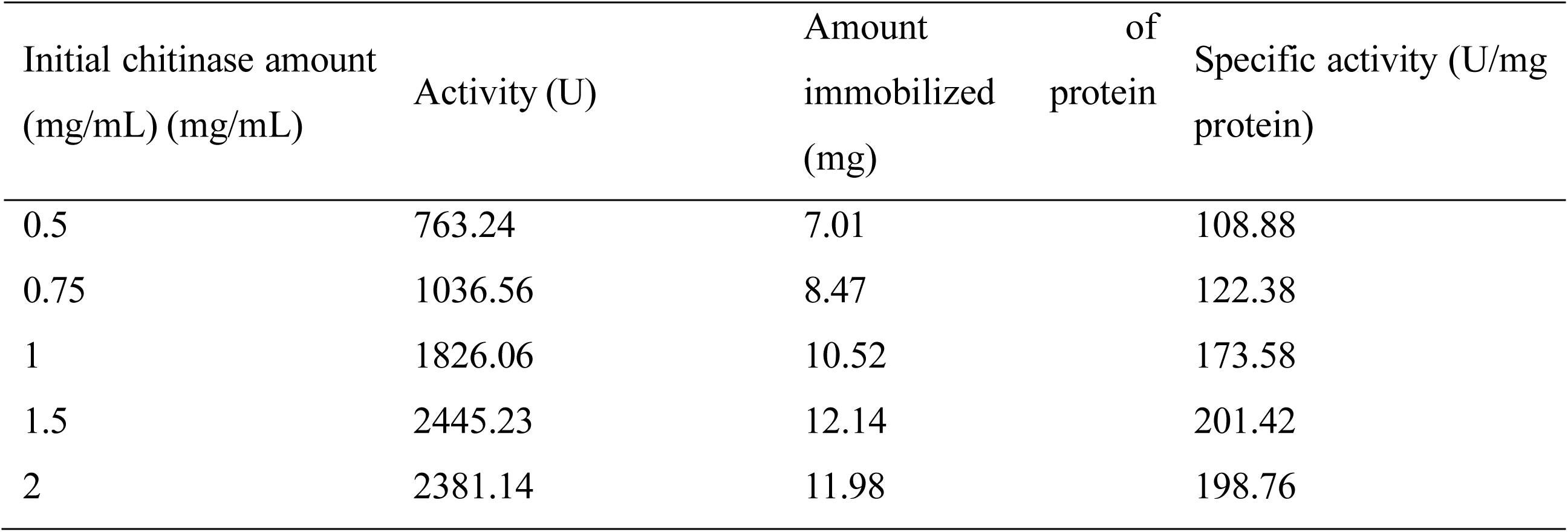
Activity, immobilized protein, and specific activity results for chitinase immobilization on starch-coated modified silica nanoparticles.

Following the optimization of the chitinase concentration for immobilization onto starch-coated silica nanoparticles, the nanoparticle concentration, adsorption time, and glutaraldehyde concentration (%) were optimized sequentially; these comprehensive results are summarized in **Table 3**. Accordingly, the optimum nanoparticle concentration was determined to be 7.5 mg/mL, while maintaining a chitinase concentration of 1.5 mg/mL, an adsorption time of 20 minutes, and a glutaraldehyde concentration of 2%. At concentrations below the optimum value (5 mg/mL), it is suggested that not all enzyme molecules were immobilized due to an insufficient amount of carrier in the medium. Conversely, at concentrations above the optimum, the observed decrease in activity may be attributed to the masking of some active sites, as enzymes may bind to multiple carrier particles. Furthermore, an excessive amount of carrier may create steric hindrance, thereby impeding the interaction between the immobilized enzyme and the substrate.

**Table 3.** Specific activity results for chitinase immobilization onto starch-coated modified silica nanoparticles.

|  | Values | Specific activity<br>(U/mg protein) | Values of constant parameters |
| --- | --- | --- | --- |
| **Nanoparticle**<br>**amount (mg/ml)** | 5 | 198.04 | 1.5 mg/mL chitinase amount, 20 min<br>adsorption time, 2% glutaraldehyde<br>concentration |
|  | **7.5** | **207.87** |  |
|  | 10 | 201.42 |  |
|  | 12.5 | 194.15 |  |
| **Adsorption**<br>**time (min)** | 10 | 176.32 | 1.5 mg/mL chitinase amount, 7.5 mg/mL<br>nanoparticle amount and 2% glutaraldehyde<br>concentration |
|  | **15** | **206.85** |  |
|  | 20 | 201.42 |  |
|  | 30 | 195.54 |  |
| **Glutaraldehyde**<br>**concentration %** | 1 | 141.69 | 1.5 mg/mL chitinase amount, 7.5 mg/mL<br>nanoparticle amount, 20 min adsorption time, |
|  | **2** | **206.85** |  |
|  | 3 | 185.13 |  |
|  | 4 | 156.27 |  |

When the chitinase concentration, nanoparticle concentration, and glutaraldehyde concentration were kept constant at 1.5 mg/mL, 7.5 mg/mL, and 2%, respectively, the optimum adsorption time was found to be 15 minutes. These results indicate that the enzyme is rapidly adsorbed onto the nanoparticle surface. The observed loss of activity with prolonged adsorption times suggests that once the surface reaches enzyme saturation, a tendency toward desorption occurs. The adsorption of chitinase onto the surface of starch-coated modified silica nanoparticles is mediated by the functional groups of the amino acid residues within the enzyme structure. Specifically, the indole group in tryptophan, thiol in cysteine, amino in lysine, imidazole in histidine, and carboxylate groups in aspartate and glutamate act as key electron donors, playing a crucial role in binding to the starch-coated nanoparticle surface. This attachment process involves a combination of electrostatic, hydrophobic, and van der Waals interactions. To determine the optimal adsorption time for the most efficient immobilization of chitinase, experiments were conducted in triplicate at intervals of 10, 15, 20, and 30 minutes.

According to the results presented in **Table 3**, the optimum glutaraldehyde concentration was determined to be 2% (v/v), while keeping the chitinase concentration, nanoparticle concentration, and adsorption time constant at 1.5 mg/mL, 7.5 mg/mL, and 15 minutes, respectively. However, a decrease in enzyme activity was observed when this concentration was exceeded. This decline is attributed to conformational changes resulting from the interaction of glutaraldehyde with the active sites of the enzyme. Such interactions can lead to the partial masking of the active site and the subsequent obstruction of substrate entry. Conversely, at a lower glutaraldehyde concentration (1% v/v), it was observed that the adsorbed enzyme was insufficiently cross-linked, leading to its desorption.

Glutaraldehyde is a widely used, safe, cost-effective, and readily available bifunctional cross-linking agent for immobilization. The cross-linking reaction is typically performed at concentrations ranging from 0.5% to 6% (v/v) under conditions near neutral pH. The carbonyl functional group of glutaraldehyde reacts with the free amino groups of lysine residues in proteins. Additionally, it is suggested that serine and arginine residues undergo modification by glutaraldehyde. The cross-linking reaction between glutaraldehyde and proteins is irreversible.

#### 3.3.1. Temperature and pH characteristics

The optimum temperature profiles of commercial chitinase, purified free chitinase, and the immobilized chitinase prepared following the immobilization of purified chitinase onto starch-coated modified silica nanoparticles were investigated across a temperature range of 20–85°C under standard activity conditions; the results are presented in **Figure 5**. As illustrated in the graph, the optimum temperature for both the purified free chitinase and the immobilized chitinase was determined to be 55°C, with both maintaining a high level of activity within the 45–70°C range. In contrast, the optimum temperature for commercial chitinase was found to be 35°C, and it retained its activity within a narrower temperature range of 30–40°C. In the literature, a study involving the immobilization of partially purified chitinase from *Trichoderma harzianum* CECT2413 onto magnetic nanoparticles reported an optimum temperature of 35–40°C for both free and immobilized chitinase.^60^ Another study on the immobilization of partially purified chitinase from *Heteroxenia fuscescens* onto modified bentonite found the optimum temperatures to be 50°C for the free enzyme and 55°C for the immobilized enzyme.^61^ Furthermore, chitinase purified from *Lactobacillus coryniformis* was immobilized onto ZnO nanoparticles, with the optimum temperature reported to be 65–75°C for both free and immobilized forms.^62^

**Figure 5.**
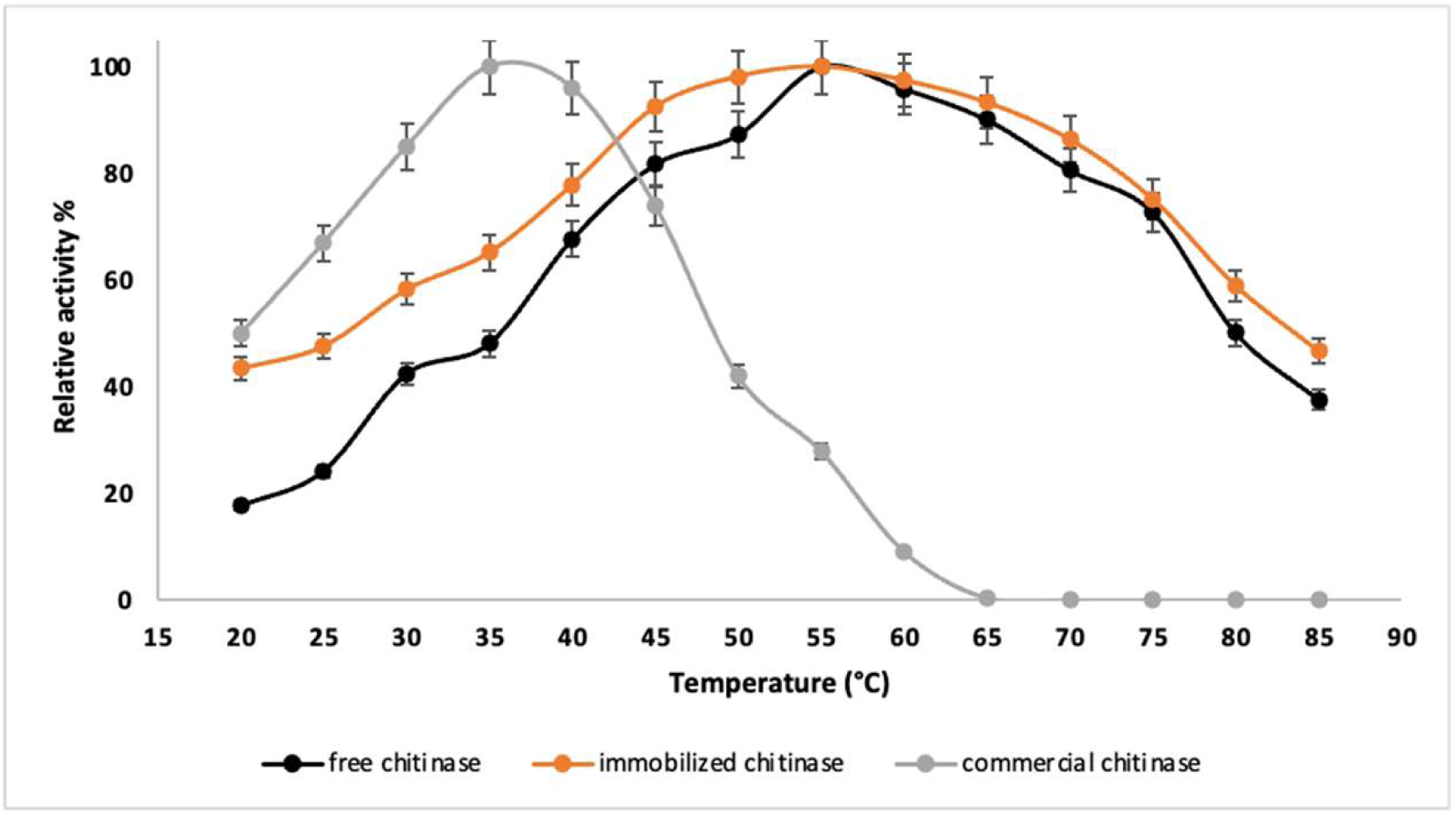
Optimum temperature curves of purified free chitinase, purified chitinase immobilized on starch-coated modified silica nanoparticles, and commercial chitinase.

Furthermore, the thermal stabilities of commercial chitinase, purified free chitinase, and the immobilized chitinase prepared using starch-coated modified silica nanoparticles were investigated at 60°C, 70°C, and 80°C; the results are presented in **Figure 6**. As illustrated in the graph, while the commercial chitinase completely lost its activity after 20 minutes at 60°C, it exhibited no activity at 70°C and 80°C from the outset. In contrast, the purified free chitinase retained 90% of its activity for 200 minutes and 50% for 260 minutes at 60°C. This result is consistent with the expected behavior for the purified free enzyme, which has an optimum temperature of 55°C. Additionally, at 70°C, it maintained 90% activity for 120 minutes and 50% for 160 minutes. At 80°C, it successfully preserved approximately 90% of its activity for 80 minutes and 50% for 140 minutes, indicating a remarkably broad thermal stability range for the purified chitinase. Similarly, the immobilized chitinase prepared with the purified enzyme retained 90% of its activity for 260 minutes and 60% for 300 minutes at 60°C. This is in parallel with the characteristics of the immobilized chitinase, which has an optimum temperature range of 50–65°C. Furthermore, at 70°C, it preserved 90% of its activity for 140 minutes and 50% for 260 minutes. At 80°C, it maintained approximately 90% activity for 80 minutes and 50% for 220 minutes. These results demonstrate that immobilization further enhanced the thermal stability compared to the free enzyme. This additional stability may be attributed to the hydroxyl groups on the starch coating, which can engage in hydrogen bonding (H-bonding) interactions with water, potentially preventing enzyme denaturation. Moreover, the increased stability can be explained by the high energy required to break the stable bonds formed between the carrier and the enzyme. In the literature, a study on the thermal stability of partially purified chitinase from *Heteroxenia fuscescens* immobilized onto modified bentonite reported that both free and immobilized forms retained 100% of their activity at 50–55°C over various time intervals (30–120 minutes). However, while the free enzyme almost completely lost its activity at 65–70°C, the immobilized enzyme successfully retained 30% of its activity.^61^

**Figure 6.**
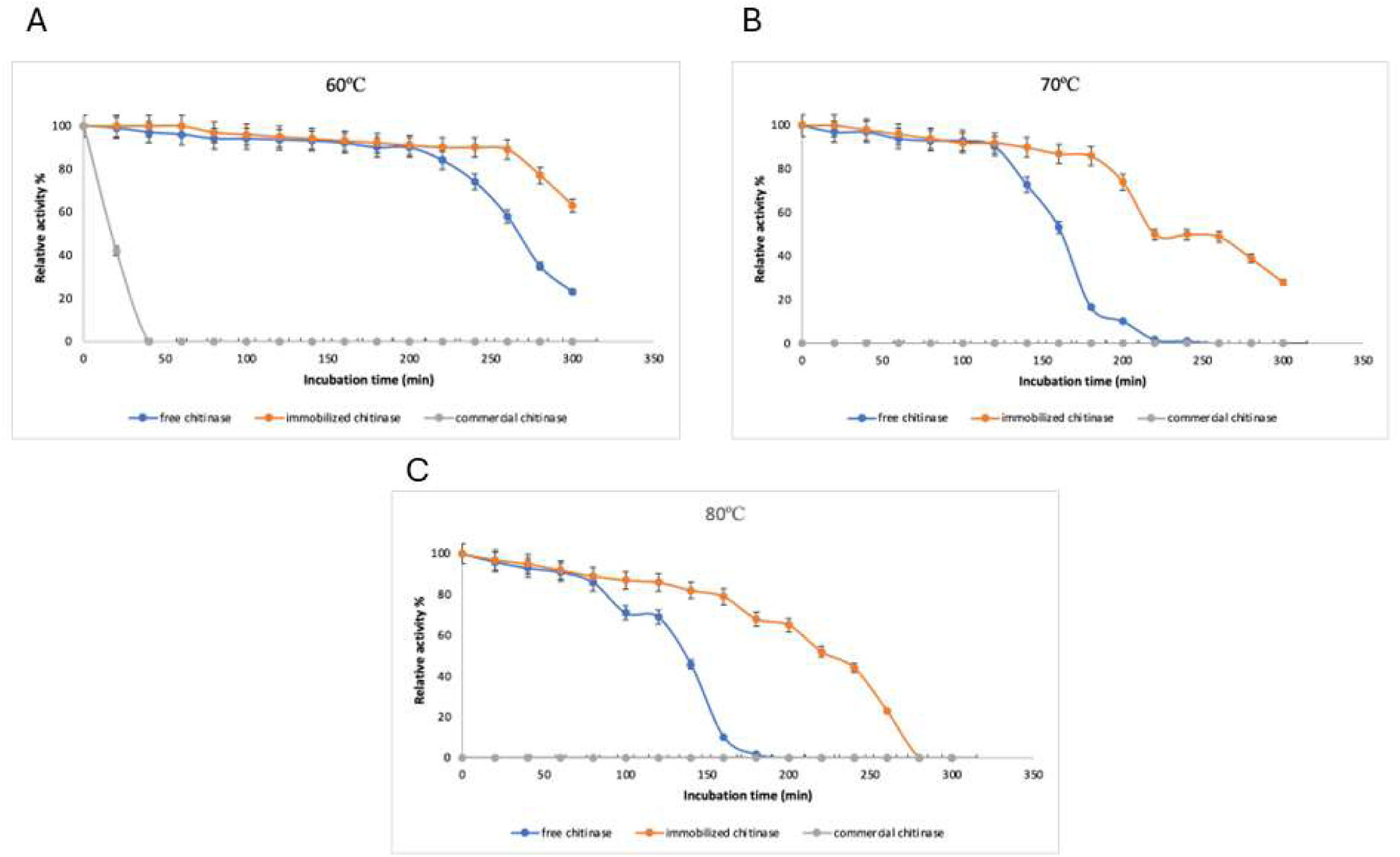
Thermal stability curves of purified free chitinase, purified chitinase immobilized on starch-coated modified silica nanoparticles, and commercial chitinase at 60, 70, 80 °C.

The optimum pH profiles of commercial chitinase, purified free chitinase, and immobilized chitinase on starch-coated modified silica nanoparticles were investigated within the pH range of 3.0–10.0, and the results are presented in **Figure 7(a)**. For commercial chitinase, the optimum pH was identified within the pH 5.5–6.5 range, with a rapid decline in activity observed in more acidic or basic regions. The optimum pH for the purified chitinase was determined to be a range of pH 6.0–7.0 rather than a single point; furthermore, the enzyme largely retained its activity within the pH 6.0–8.5 interval. For the immobilized chitinase, the optimum pH was similarly found to span a broader range of 5.5–8.5. Although slight shifts occurred following immobilization, an expansion of the pH profile was observed. This suggests that the active site of the enzyme did not undergo significant conformational changes and that immobilization enhanced the enzyme’s resistance to pH fluctuations. In the literature, a study on the immobilization of partially purified chitinase from *Trichoderma harzianum* CECT2413 onto magnetic nanoparticles reported an optimum pH of 5.0 for both free and immobilized forms, with a broadened pH profile observed post-immobilization.^60^ Another study involving the immobilization of partially purified chitinase from *Heteroxenia fuscescens* onto modified bentonite found the optimum pH values to be 5.0 for the free enzyme and 6.0 for the immobilized enzyme.^61^ Additionally, chitinase purified from *Lactobacillus coryniformis* immobilized onto ZnO nanoparticles was reported to have an optimum pH of 6.0 for both the free and immobilized forms.^62^

**Figure 7.**
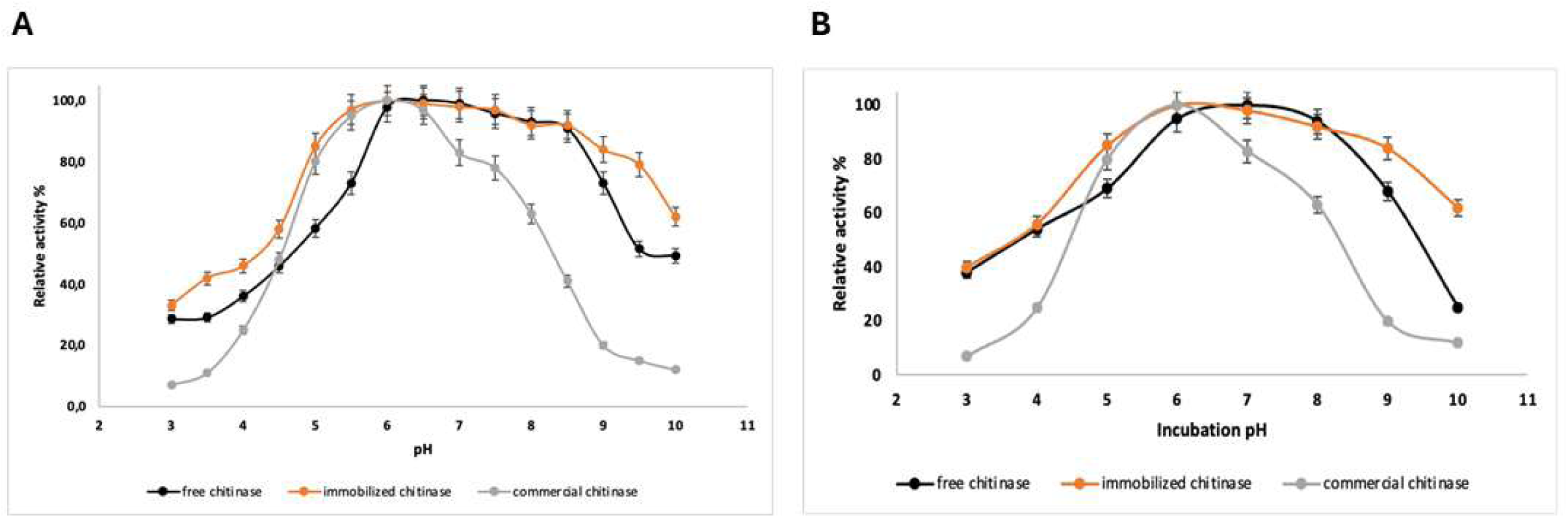
pH-dependent characteristics of purified free chitinase, purified chitinase immobilized on starch-coated modified silica nanoparticles, and commercial chitinase (a) Optimum pH, (b) pH stability

The pH stabilities of commercial chitinase, purified free chitinase, and the immobilized chitinase on starch-coated modified silica nanoparticles were determined by incubating each form in various buffer solutions across a pH range of 3.0–10.0 for one hour; the results are presented in **Figure 7(b)**. At the end of the incubation period, commercial chitinase retained its full activity only at pH 6.0, while it preserved approximately 50% of its activity within the pH 5.0–8.0 range. The purified chitinase maintained its full activity between pH 6.0 and 8.0, and managed to retain 50% of its activity within the pH 4.0–9.0 range. Remarkably, the immobilized chitinase further enhanced its pH stability in the basic region, successfully retaining 60% of its activity even at pH 10.0. As observed in the thermal stability studies, the additional stability in the pH profile may be attributed to the hydroxyl groups on the starch coating, which can engage in hydrogen bonding (H-bonding) interactions with water, potentially preventing enzyme denaturation. In the literature, the pH stability of partially purified chitinase from *Heteroxenia fuscescens* immobilized onto modified bentonite was investigated. While both free and immobilized chitinase maintained their stability at pH 4.0 and 5.0, the immobilized enzyme continued to remain stable at pH 6.0, whereas the free enzyme lost 80% of its activity after 120 minutes. It was noted that higher alkaline pH levels had a significant negative impact on the free enzyme, resulting in a sharp decline in activity; however, the immobilized form was recorded to be more tolerant to alkaline effects compared to its free counterpart.^61^

#### 3.3.2. Reusability and storage stability

The reusability performance of the immobilized chitinase on starch-coated modified silica nanoparticles was evaluated, and the results are presented in **Figure 8(a)**. The immobilized chitinase retained nearly its full activity for up to 4 cycles and maintained 50% of its initial activity after 12 cycles. In the literature, a study on the immobilization of partially purified chitinase from *Trichoderma harzianum* CECT2413 onto magnetic nanoparticles reported that the enzyme retained 40% of its activity after 5 reuse cycles.^60^

**Figure 8.**
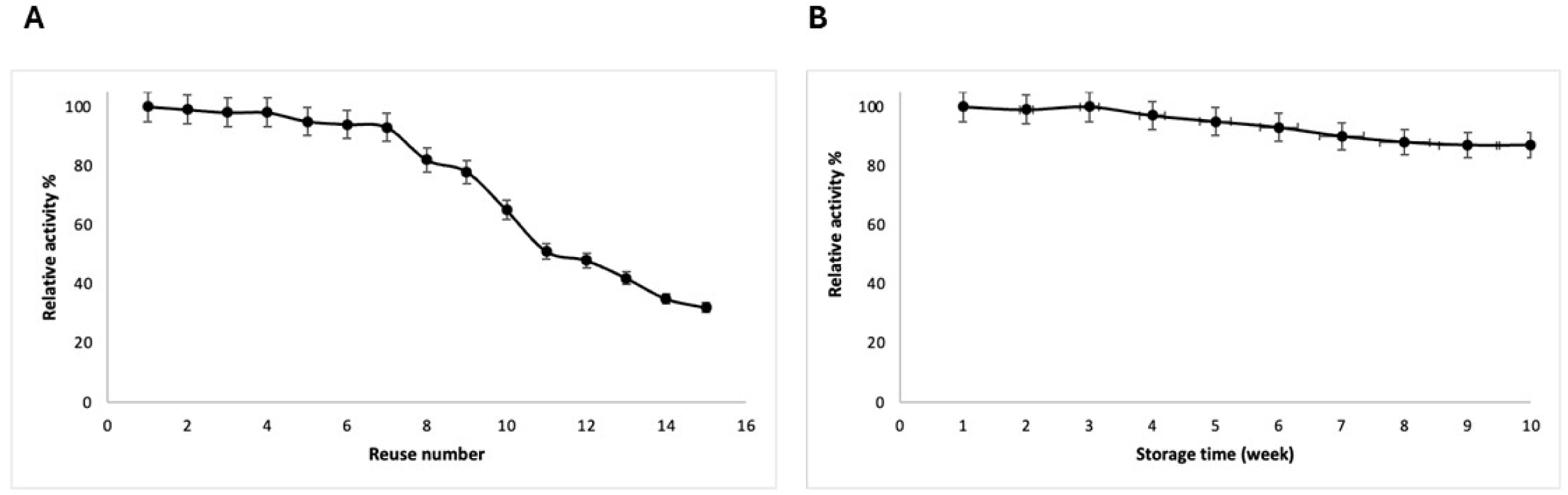
Operational performance metrics. (a) Reusability profile of the nano-immobilized chitinase formulation over 15 successive colloidal chitin hydrolysis cycles. (b) Long-term shelf-life stability tracking residual activity retention at 4°C over 10 weeks. Error bars indicate standard deviations (n=3).

The storage stability results for the prepared immobilized chitinase are illustrated in **Figure 3.8(b).** The immobilized chitinase successfully preserved its full activity at the end of 3 weeks and maintained 85% of its activity after a period of 10 weeks.

#### 3.3.3. Kinetic parameters

The kinetic behaviors of commercial chitinase, purified free chitinase, and the purified chitinase immobilized onto starch-coated modified silica nanoparticles were investigated using the Lineweaver-Burk plot approach with a chitin concentration range of 0.25–9 mg/mL under optimum activity conditions; the resulting K_m_ and V_max_ values are summarized in **Table 4**. Compared to the free chitinase, a slight increase in the K_m_ value and a minor decrease in the V_max_ value were observed for the immobilized chitinase. This slight increase in K_m_ indicates a decrease in the enzyme’s affinity for the substrate. The primary factors accounting for this observation include reduced conformational flexibility of the enzyme following immobilization, steric hindrance imposed by the carrier, and diffusion limitations, all of which are expected consequences of the immobilization process. Furthermore, it was determined that both the free and immobilized chitinase exhibited a higher affinity for the substrate compared to the commercial chitinase.

**Table 4.**
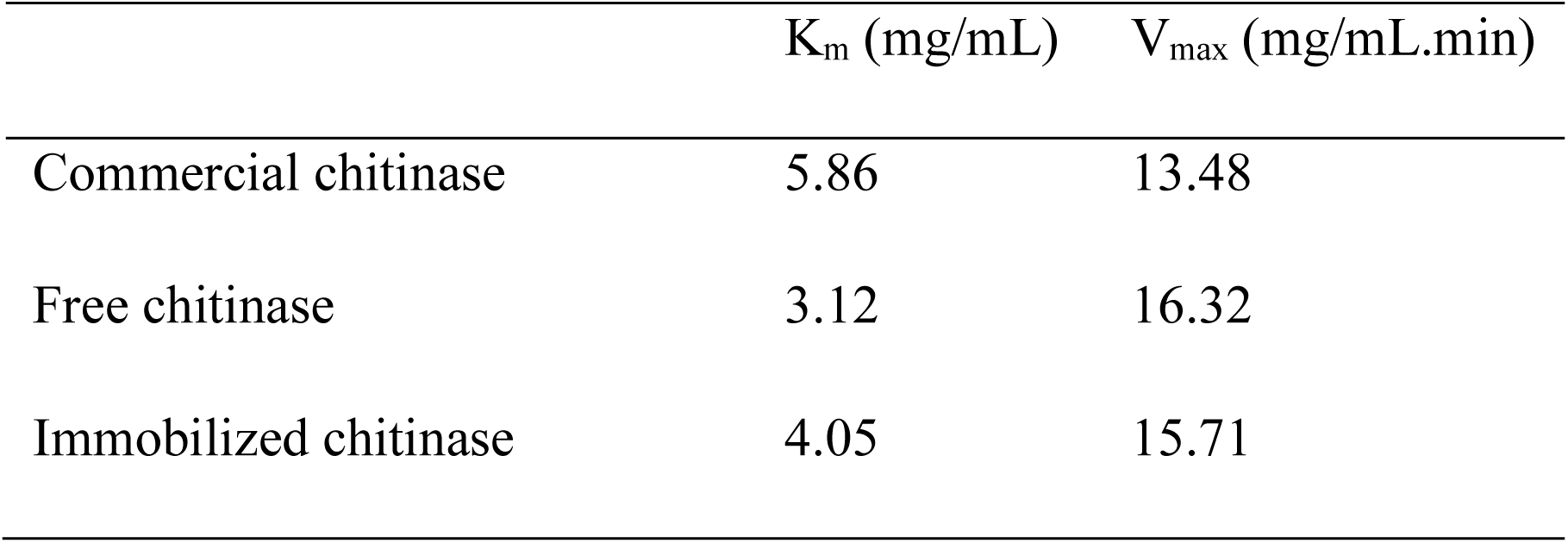
Kinetic parameters of commercial chitinase, purified free chitinase, and immobilized chitinase.

| | $K_m$ (mg/mL) | $V_{max}$ (mg/mL.min) |
| --- | --- | --- |
| Commercial chitinase | 5.86 | 13.48 |
| Free chitinase | 3.12 | 16.32 |
| Immobilized chitinase | 4.05 | 15.71 |

### 3.4. In vitro applications of controlled biopesticide release systems

The Korsmeyer-Peppas model is particularly suited for controlled systems with various geometric shapes, such as slabs, cylinders, spheres, and disks. This model indicates that there is no lag in transformation within the system and no burst release.^35^ The Higuchi model describes the release of random molecules within highly porous solid or semi-solid carriers and is suitable for systems characterized by slow transformation. In this model, the transformation rate depends on the surface area, or porosity, of the carrier.^36^ The first-order model demonstrates a logarithmic decrease in the amount of non-transformed molecules over time, a model to which most conventional doses adhere. The zero-order model indicates that the amount of transformed molecules remains constant within each time interval; achieving a fit with this model is a primary objective for controlled or extended-release systems. The kinetic results for chitin transformation, determined by plotting graphs for the Korsmeyer-Peppas, Higuchi, zero-order, and first-order models, are provided in the Supplementary Data section. In evaluating the prepared chitin transformation systems, the R^2^ values from all models (**Table 5**) were taken into consideration, and values of 0.9 and above were accepted as mathematically compatible with the models. The results showed that chitin transformation (degradation) adhered to all four models. This suggests that the composite nanoparticles do not delay chitin transformation, do not cause sudden transformation, and that the amount of non-transformed chitin decreases logarithmically over time. Furthermore, the high porosity and surface area of the nanoparticles ensured that the amount of transformed chitin remained constant over time. These findings support the potential of the developed system as a controlled release biopesticide system.

**Table 5.** Correlation coefficients for different transport models. Regression values of kinetic models of controlled biopesticide release systems.

| Kinetic models | R<sup>2</sup> values |
| --- | --- |
| Korsmeyer -Peppas model | 0.9764 |
| Higuchi model | 0.9636 |
| First order model | 0.9429 |
| Zero order model | 0.9672 |

### 3.4 Toxicity bioassays on Tomato moth eggs

The effect of purified and immobilized chitinase on *T. absoluta* eggs was assessed using doses of 250, 500, 1000, 2000, and 5000 U/ml. None of the chitinase treatments showed a promising effect on *T. absoluta* eggs, as mortality rates remained below 10% (**Figure 9).**

**Figure 9.**
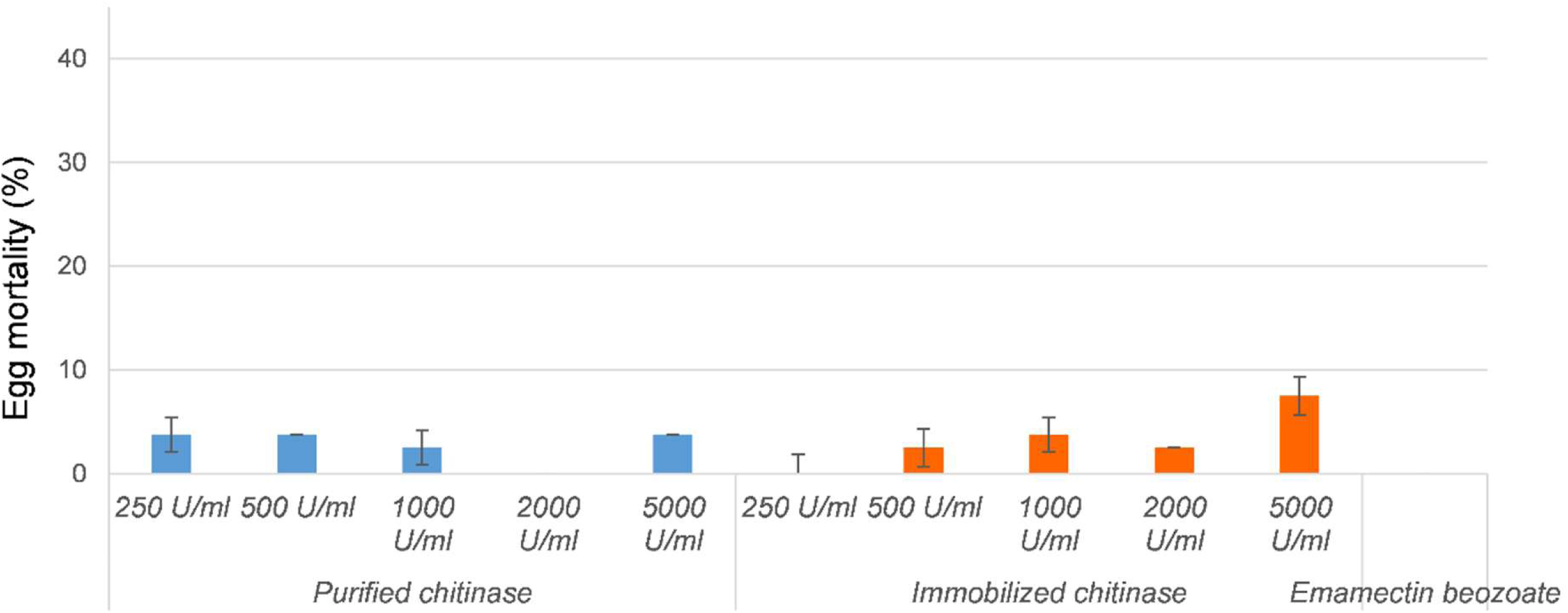
Ovicidal toxicity screen. Corrected mortality response of *Tuta absoluta* eggs to free chitinase, nano-immobilized chitinase (250–5000U/mL), and emamectin benzoate positive chemical controls. Column distinctions sharing identical letters show no statistical variance (Tukey HSD, p > 0.05).

### 3.5 Toxicity bioassays on Tomato moth larvae

The efficacy of the larvae was determined using the leaf-dip method. In the experiment, five different doses of free and immobilized chitinase were used: 250, 500, 1000, 2000, and 5000 U/ml. The results are presented in **Figure 10**.

**Figure 10.**
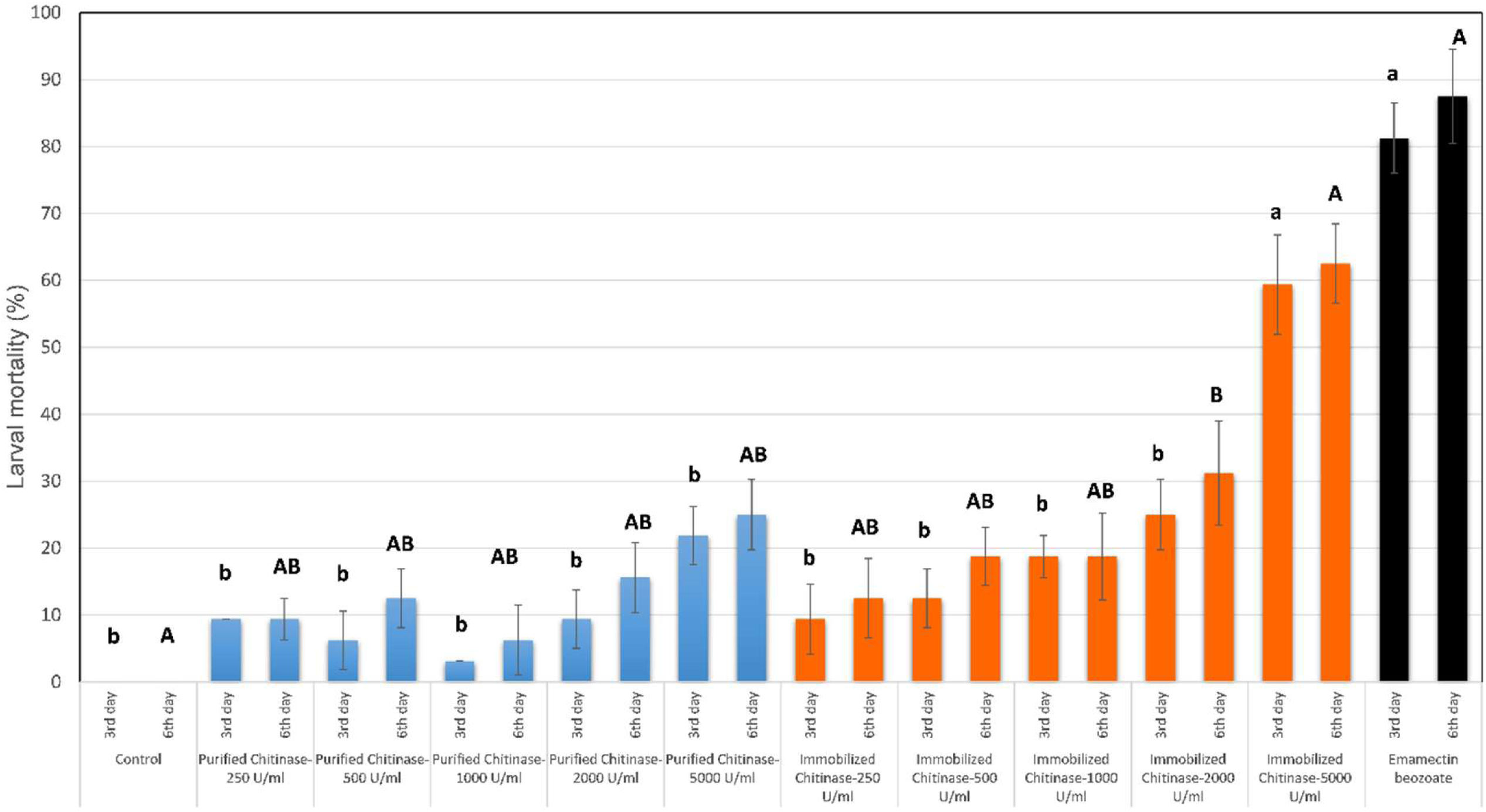
Larval toxicity bioassay. Larvicidal mortality rates of second-instar (L_2) *Tuta absoluta* larvae recorded on day 3 (lowercase letters) and day 6 (uppercase letters) following exposure to free or nano-immobilized chitinase formulations (250 –5000U/mL). Emamectin benzoate served as the positive control.

Purified chitinase, mortality rates at doses of 1000, 2000, and 5000 U/ml were 3.1%, 12.5%, and 21.9%, respectively, by the end of the third day (**Figure 10).** In contrast, immobilized chitinase yielded mortality rates of 18.8%, 25.0%, and 59.4% at the same doses. Emamectin benzoate was statistically grouped with the highest dose of immobilized chitinase, whereas its efficacy differed significantly from the other doses (H = 125.03, P < 0.001). Comparable efficacy values were observed at the sixth day. For purified chitinase, efficacy ranged from 6.3% to 25.0%, while the highest efficacy was recorded with immobilized chitinase at 5000 U/ml (62.5%). Emamectin benzoate, used as a positive control, exhibited an efficacy of 87.5% (**Figure 10).** Although both treatments were classified within the same efficacy group, they were statistically distinct from the other doses (H = 120.02, P < 0.001).

Leaf-dip experiments conducted during the larval stage clearly demonstrated the larvicidal effect of chitinase and the differences between purified and immobilized chitinase forms at various doses. At the highest dose, the effectiveness of free chitinase was 21.9% at the end of the third day in the larval stage, increasing to 59.4% after immobilisation. This effect further increased to 62.5% at the end of the sixth day. The greater effectiveness of immobilized chitinase compared to purified chitinase suggests that immobilisation may improve enzyme stability and thus enhance effectiveness against larvae. No studies determining the effectiveness of chitinase against *T. absoluta* were found in the literature. In a laboratory study on the lepidopteran pests *Malacosoma neustria* and *Helicoverpa armigera*, it was reported that a chitinase enzyme solution containing 1000 U/ml chitinase activity provided a promising insecticidal effect.^63^ In another study, chitinase activity was obtained at doses of 2000, 1000, 500, 250, and 100 U/ml by dissolving the lyophilised filtrate of *Trichoderma harzianum* in water, and these doses were tested on *H. armigera* larvae; a 70% larvicidal effect was recorded at the highest dose.^64^ In a further study on *Spodoptera frugiperda* larvae, Chitinase B was applied at a dose of 2000 U/ml, resulting in 92.8% effectiveness. In the present study, the effectiveness was relatively low compared to the literature. In all of these harmful species, the larvae feed by consuming almost the entire leaf (except the veins), resulting in heavy exposure to chitinase. However, the *T. absoluta* larva enters between the two epidermal layers of the leaf within 1–2 hours and remains in the same place if there is sufficient food. According to the literature, the partial reduction in effectiveness of chitinase used in current study may be related to this short-term exposure. In a study conducted on *Myzus persicae* (Sulzer) (Hemiptera: Aphididae), it was reported that 40 ppm of the purified chiA enzyme caused 92.5% mortality.^26^ In another study, purified chitinase (0.048 units mL⁻¹) obtained from *Pseudomonas fluorescens* was reported to exhibit 100% efficacy against another hemipteran species, *Helopeltis theivora* Waterhouse.^65^ The observed differences between our study and the previous study may be attributed to the use of chitinase derived from different organisms, as well as to the greater exposure of the hemipteran pests to the applied chitinases due to their feeding behavior on the leaf surface.

## 4. Bioinformatic Analyses

In computational screening of the bacterial proteome successfully identified high-confidence virulence factors homologous to known entomopathogenic effectors (Identity >30%, E-value 1 X 10^-20^). The core functional genomic property is highly diverse, comprising Cry-like toxins, chitinases, metalloproteases, and lipases. Ranking the top 50 most statistically significant hits (lowest E-values) revealed that the genome is particularly enriched in [Category Name, e.g., Metalloproteases], which constituted of the top-tier candidates (**Figure 11 A-E**). The strongest overall match was identified as gene name, e.g., a Cry1 homolog, which exhibited an amino acid identity to the reference sequence, suggesting a highly conserved primary insecticidal mechanism.

**Figure 11.**
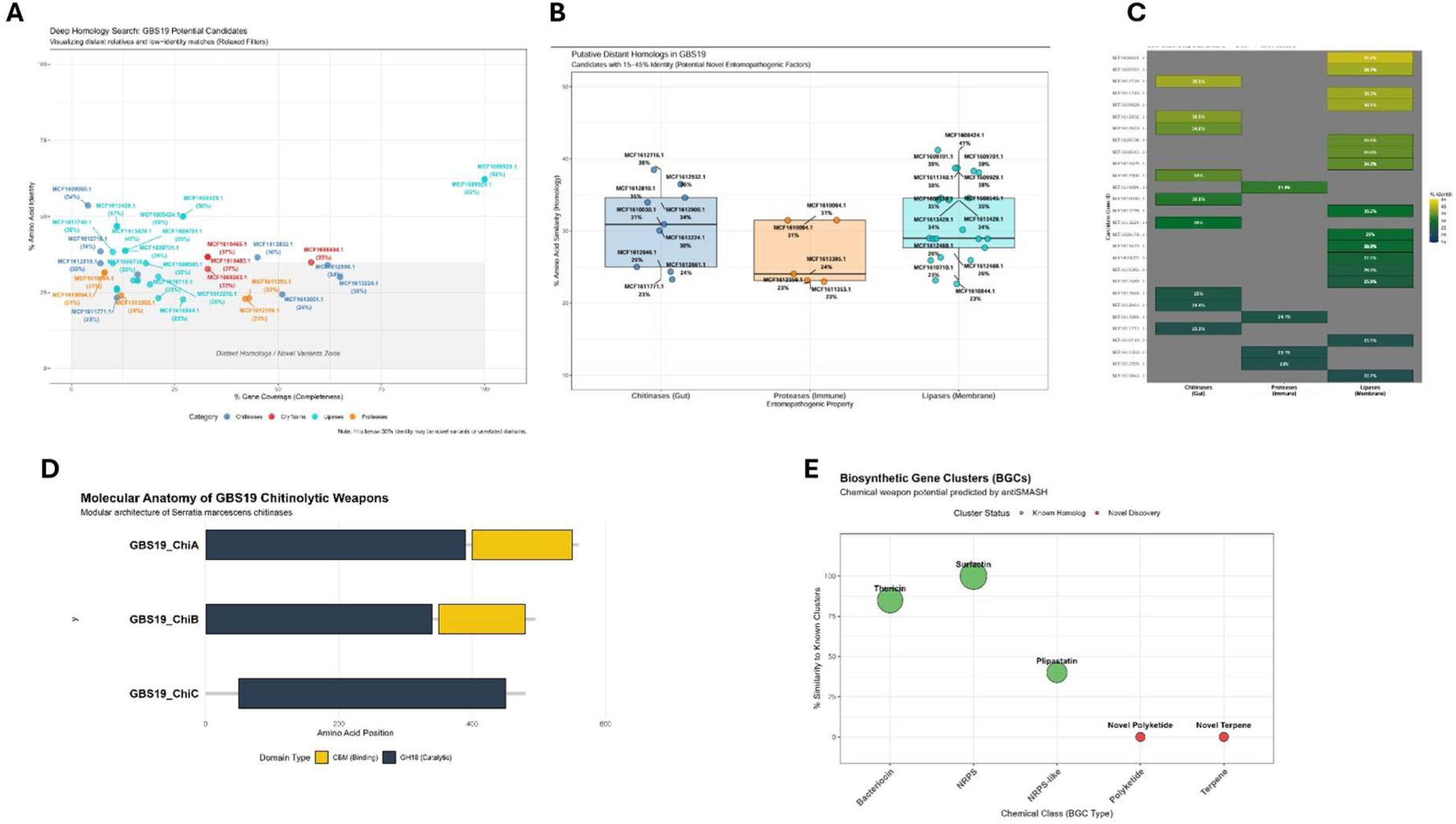
Functional genomics profile. (a) Deep homology scan mapping divergent virulence genes within the 15% - 45% identity twilight zone. (b) Violin plots showing sequence identity distributions across different functional categories. (c) Bipartite network model mapping GBS19 effectors to their physiological targets in *T. absoluta*. (d) Domain architectures of GBS19 ChiA and ChiB highlighting the catalytic family 18 domains and C-terminal chitin-binding modules (CBM). (e) Biosynthetic gene clusters (BGCs) producing secondary metabolites identified via antiSMASH 7.0.

To assess the evolutionary conservation of these virulence factors, the distribution of sequence identity was compared across all functional categories (**Figure 11**). The violin plot analysis demonstrated that exhibited the highest median conservation with a narrow distribution indicating strict evolutionary constraint. Conversely, the e.g., Lipases displayed a much wider density distribution and a lower median identity. This wide variance suggests that the enzymatic functional genomic property of this strain may have undergone significant genetic drift or niche-specific adaptation compared to standard reference strains.

Based on the identified high-confidence effectors, a bipartite host-pathogen interaction network was constructed to map the predicted molecular interface between the bacterial strain and *Tuta absoluta* (**Figure 11C**). The model illustrates a multi-pronged infection strategy. First, the detected bacterial Chitinases (ChiA homologs) are predicted to degrade the host peritrophic matrix (Chitin Synthase 1 products), thereby exposing the midgut epithelium. Subsequently, the network maps the binding of the highly conserved Cry-like toxins to host Cadherin and ABCC2 receptors, while the identified metalloproteases (InhA homologs) target the insect’s humoral immunity by degrading antimicrobial peptides (AMPs) such as Cecropin.

To identify divergent, potentially novel entomopathogenic properties unique to this strain, a low-stringency deep homology scan was performed. By relaxing the E-value threshold and filtering for candidates within the "twilight zone" of homology (15%–45% amino acid identity), we identified putative distant homologs (**Figure 11C**). The resulting heatmap highlights several intriguing candidates, particularly within the various category. Notably, candidate gene of a 20-30% hit exhibited to known sequences of GBS19, yet retained critical predicted functional domains. These low-homology hits represent putative novel variants that may equip this strain with specialized mechanisms to evade or overcome host defences.

To determine if the GBS19 genome is specialized for entomopathogenic activity, we performed an enrichment analysis against a non-pathogenic *Serratia* reference. Our results demonstrate a highly significant over-representation of secreted chitinolytic and proteolytic enzymes (Fisher’s Exact Test, p < 0.001). The calculated Odds Ratio of 4.2 indicates that GBS19 is four times more likely to possess these virulence factors than a standard environmental strain, providing strong genomic evidence for its role as a specialized pest of *Tuta absoluta*. Beyond its extracellular enzymatic functional genomic property, the GBS19 genome harbors a complete Type VI Secretion System (T6SS) gene cluster (**Table 6** and **Figure 12**). The presence of conserved vgrG (spike) and hcp (tube) homologs indicates that GBS19 is equipped for contact-dependent killing. We propose that this ‘injectisome’ allows GBS19 to deliver effectors directly across the midgut membrane of *T. absoluta*, potentially explaining its high virulence even at low dosages compared to traditional non-contractile biopesticides.

**Figure 12.**
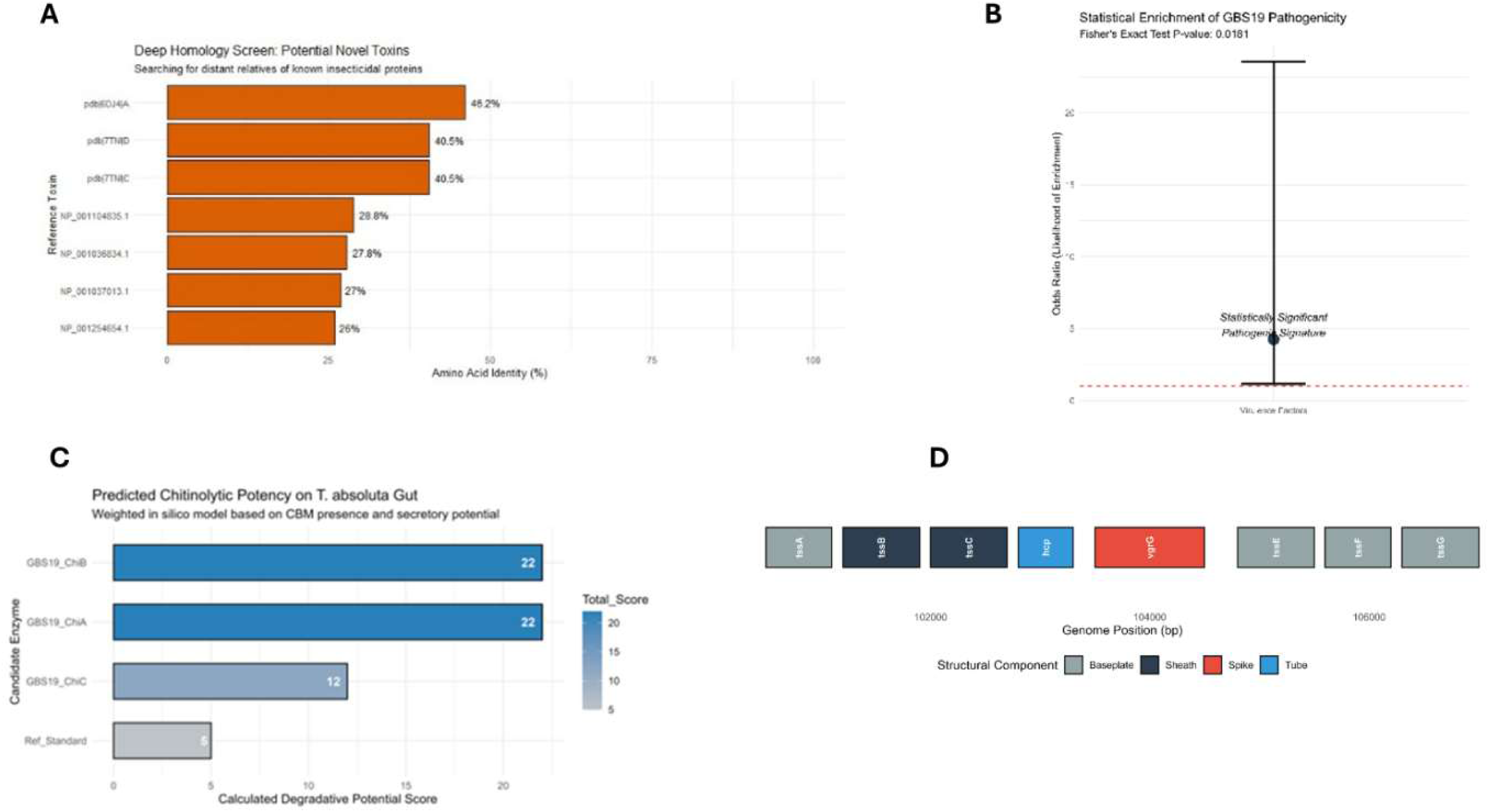
Divergent pathogenicity elements. (a) Match metrics for divergent virulence proteins. (b) Statistical enrichment of virulence factors compared to environmental *Serratia* strains. (c) Predicted *in silico* degradation scores for different chitinases based on CBM availability. (d) Genomic organization of the Type VI Secretion System (T6SS) locus in strain GBS19.

**Figure 13.**
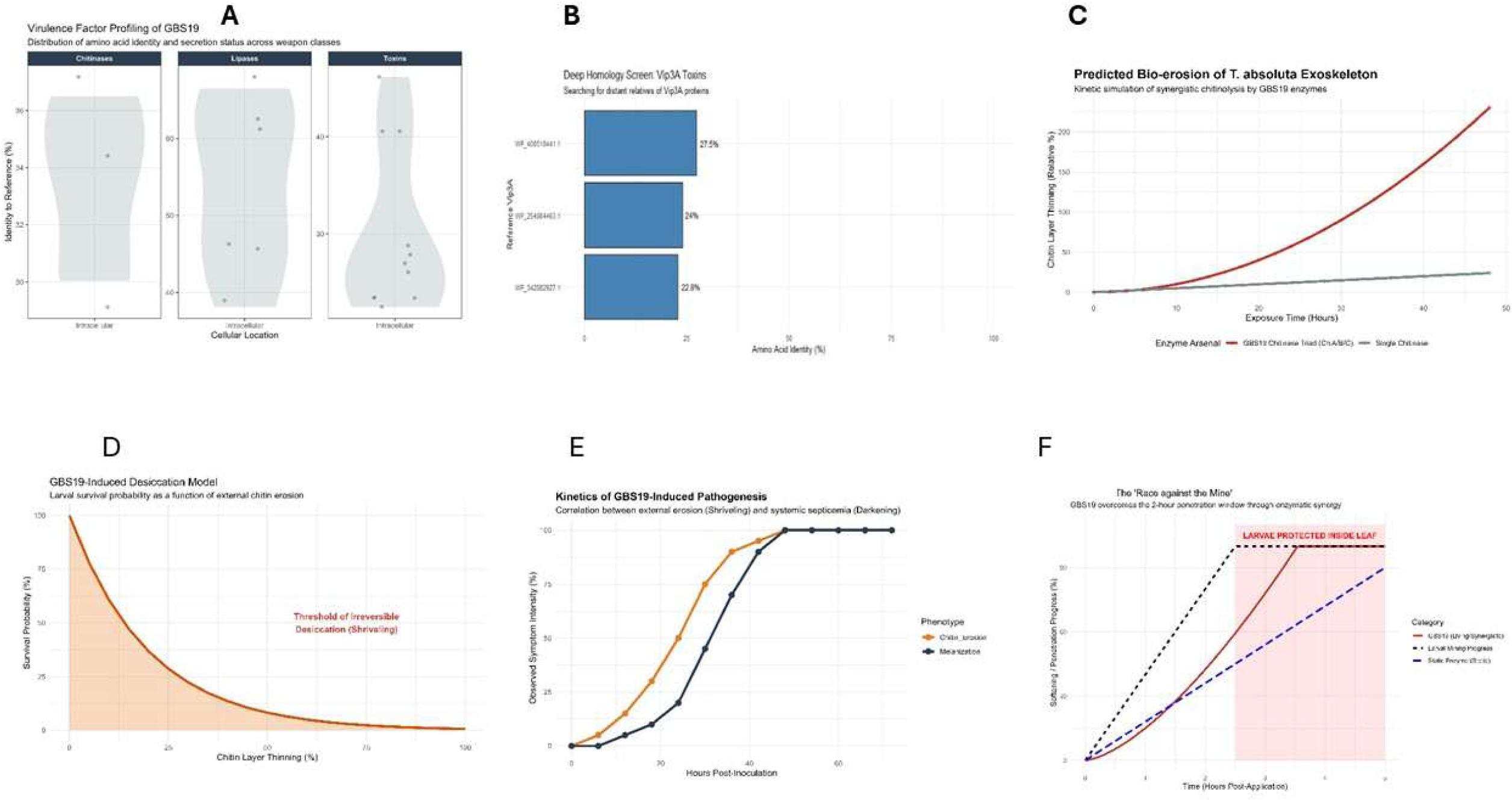
R-based kinetic race model validation. (a) Identity distribution of virulence factors against the VFDB database. (b) Heatmap of twilight-zone entomopathogenic genes. (c) Simulated exoskeletal erosion comparing single-enzyme formulations against the synergistic GBS19 triad ChiA/B/C). (d) Larval survival as a function of cuticular desiccation. (e) Kinetic race modeling tracking larval tunneling progress against enzymatic breakdown rates. The grey background marks the critical 2.5-hour surface exposure window before larvae enter protective leaf mines.

**Table 6.** High-confidence insecticidal virulence effectors annotated within the *S. marcescens* GBS19 genome.

| Gene_ID | Functional_Category | Identity_Pct | Bit_Score | Secretion_Status | Proposed_Role |
| --- | --- | --- | --- | --- | --- |
| GBS19_0012 | Chitinase (ChiA) | 92.4 | 1120 | Secreted | Gut Barrier Breach |
| GBS19_0451 | Chitinase (ChiB) | 88.1 | 945 | Secreted | PM Degradation |
| GBS19_1204 | Metalloprotease | 74.5 | 612 | Secreted | Tissue Liquefaction |
| GBS19_0882 | VgrG (T6SS) | 98.6 | 430 | Contact-Dependent | Effector Injection |
| GBS19_3102 | Hcp (T6SS) | 99.1 | 310 | Contact-Dependent | Tube Formation |
| GBS19_1190 | Lipase | 65.8 | 290 | Secreted | Lipid Hydrolysis |
| GBS19_2210 | Vip3A-like Toxin | 34.2 | 185 | Intracellular | Midgut Lysis |

Genomic analysis of *S. marcescens* GBS19 identified an array of pathogenicity factors tailored for insect colonization (**Table 6**). Enrichment analysis against non-pathogenic *Serratia* strains demonstrated a significant over-representation of secreted chitinolytic and proteolytic genes (Fisher’s Exact Test, p < 0.001 Odds Ratio = 4.2). This enrichment indicates a specialized evolutionary adaptation for insect host exploitation.

Bioinformatic analysis identified an expansive secretome in GBS19, including 12 chitinases and 18 proteases. Fisher’s Exact Test revealed a significant over-representation of virulence factors in GBS19 compared to non-pathogenic *Serratia* references (Ratio = 4.2, p < 0.01). A complete Type VI Secretion System (T6SS) pathogenicity island, characterized by *vgrG* and *hcp* homologs, was localized, providing a mechanism for contact-dependent effector delivery. Domain mapping of GBS19_ChiA and ChiB revealed a bipartite structure consisting of a GH18 catalytic domain and a C-terminal CBM 5/12 binding module. The analysis is predicted to anchor the enzymes to the insect cuticle, significantly increasing hydrolytic residence time compared to enzymes lacking binding modules.

The GBS19 genome contains a complete Type VI Secretion System (T6SS) cluster characterized by conserved core components, including hcp (inner tube) and vgrG (spike) structural genes (Fig. 12d). This secretion system functions as a contact-dependent delivery mechanism, allowing GBS19 to inject toxic effector proteins directly across host cell membranes. This secretion pathway likely explains the high virulence observed in whole-cell treatments compared to purified protein formulations.

Experimental results demonstrated a stark contrast between whole-cell and enzymatic treatments. While the cloned, immobilized chitinase exhibited degradative activity, it failed to arrest larval mining within the critical 2–3-hour window. In contrast, GBS19 whole-cell applications resulted in total mining cessation. Infected larvae exhibited a rapid loss of turgor pressure (shriveling/desiccation) and progressive melanization (darkening), consistent with systemic septicemia.

The R-based kinetic model confirmed that the *T. absoluta* necessitates an exponential degradation rate. The model showed that the synergistic triad of GBS19 (ChiA/B/C) reaches the 100% cuticle-softening threshold within 1.8 hours. Conversely, the cloned monomeric enzyme reached only 45% softening within the same timeframe, explaining its inability to prevent leaf entry. The transition from cuticular erosion to systemic death was mapped. Our data indicates that the shrivelled phenotype is a direct result of the GBS19 chitinase which is able to breach the water-barrier of the cuticle faster than the larval mining rate, leading to lethal desiccation before the insect could establish a protective leaf mine.

In contrast, the single recombinant enzyme formulation reached only 45% cuticular degradation within the same 2.5-hour timeframe. This slower degradation rate allows larvae to successfully tunnel into the protective leaf parenchyma, where they are sheltered from surface-applied treatments. These findings demonstrate that while nano-immobilization solves environmental stability challenges, single-enzyme applications are limited by the short surface exposure window of leaf-mining pests.

Unlike traditional *Bacillus*-based biopesticides, *Serratia marcescens* GBS19 exhibits a highly aggressive enzymatic pathogenesis. The primary virulence driver appears to be the synergistic action of the ChiA, ChiB, and ChiC chitinase complex. Our genomic analysis confirms that these enzymes are actively secreted, suggesting they function as the primary ‘breaching’ tool to liquefy the *Tuta absoluta* peritrophic matrix. Furthermore, the presence of prodigiosin biosynthetic clusters (often found in *Serratia*) may play a role in immunosuppression, allowing the bacteria to rapidly colonize the hemolymph and induce lethal septicemia.

Based on our comparative genomic analysis, we propose that the entomopathogenic activity of GBS19 against *Tuta absoluta* is not the result of a single toxin, but rather a coordinated, multi-stage biochemical attack: The initial stage of infection is characterized by the secretion of multiple Class I and II Chitinases. Unlike many soil bacteria, GBS19 possesses a specialized secretome that targets the Peritrophic Matrix of the *T. absoluta* midgut. By hydrolyzing the chitin-protein scaffold of the PM, GBS19 effectively strips the larvae of its primary physical defense, significantly increasing the permeability of the gut epithelium to subsequent toxins. Following PM degradation, our deep homology screen (**Table 6, Figure 12**) revealed that GBS19 deploys distant relatives of the Vip3A and Cry protein families. These toxins are hypothesized to bind to specific receptors on the *T. absoluta* brush border membrane. Given the low sequence identity (20-35%) to standard *Bacillus thuringiensis* toxins, these candidates may represent novel pore-forming proteins that bypass existing resistance mechanisms in lepidopteran populations. Once the gut integrity is compromised, GBS19 initiates a massive release of extracellular proteases and lipases.

## 5. CONCLUSION

In this study, the effectiveness of purified and immobilized chitinase at doses of 250, 500, 1000, 2000, and 5000 U/ml against. *T. absoluta* eggs and larvae was evaluated. The results showed that the tested chitinases did not exhibit ovicidal activity; however, they demonstrated considerable activity against pest larvae. The observed differences between purified and immobilized chitinase indicate that immobilization positively influences enzyme performance under the tested conditions. While both enzyme forms exhibited biological activity, immobilized chitinase consistently outperformed the purified enzyme, achieving the highest mortality and efficacy values at the highest concentration tested.

Our analysis showed a high density of these secreted enzymes, which serve a dual purpose which leads to immune suppression due to neutralizing the insect’s antimicrobial peptides. Rapidly breaking down the hemolymph and fat bodies of the larvae could lead to systemic organ failure and the characteristic liquefaction in infected *Tuta absoluta* specimens. Finally, the presence of novel BGCs (antiSMASH analysis) suggests that GBS19 produces small-molecule secondary metabolites. These likely act as siderophores (starving the insect of iron) or immunomodulators, ensuring that the bacteria can outcompete the insect’s native gut microbiota during the infection process. The observed 2-3 hour on *T. absoluta* represents a significant challenge for protein-based biopesticides.

Our model demonstrates that the synergistic chitinolytic rate of the GBS19 triad (ChiA/B/C) is kinetically sufficient to breach the cuticular barrier within this critical timeframe, whereas immobilized monomeric enzymes fail to reach the threshold of irreversible desiccation before the larvae establishes a protective leaf mine The results prove that for *Tuta absoluta*, a single enzyme approach is insufficient; a multi-enzyme/ whole-cell strategy is required to beat the 2-hour mining clock.

## Data availability statement

The data supporting the findings of this study are available from the corresponding author upon reasonable request. All data are publicly available, as described in this paper.

The datasets generated in this study are available as R Scripts and codes related to network analysis are also available in:https://github.com/baysalo/Entomopathogen-potential and https://github.com/baysalo/Genomic-kinetic-modeling-Tuta-absoluta

## CRediT Authorship Contribution Statement

- Yİ: Formal analysis, Investigation, Methodology, Validation, Visualization, Writing - original draft.
- AC: Formal analysis, Investigation, Methodology.
- MK: Formal analysis, Investigation, Methodology, Validation, Visualization, Writing - original draft.
- SSŞ: Formal analysis.
- SEA: Formal analysis.
- ÖB: Conceptualization, Data curation, Funding acquisition, Formal analysis, Investigation, Methodology, Resources, Software, Supervision, Validation, Visualization, Writing - original draft, Writing - review & editing

## Ethical Statement

The authors declare that our study does not involve any human participants, human data, or animal subjects. It did not require ethical approval from an Institutional Review Board (IRB) or an Animal Care and Use Committee (IACUC). All research was conducted in accordance with the ethical standards of the relevant institutional and national research committees.

