## Supplementaryfile for "Genomic and Kinetic Modeling Involving Nanoparticle-Mediated Delivery of a Novel Chitinase Enzyme to Outpace *Tuta absoluta* Damage"

**Supplementary Data**

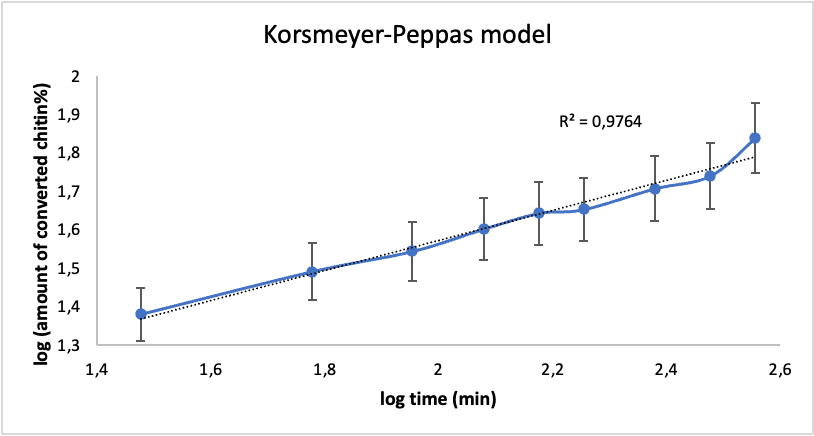

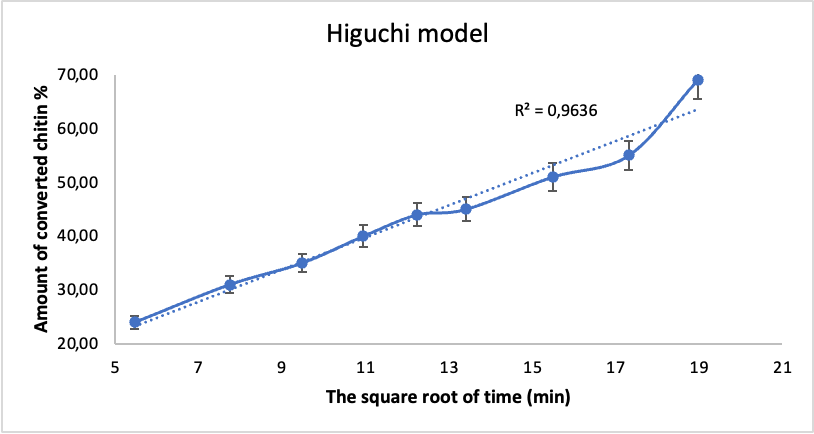

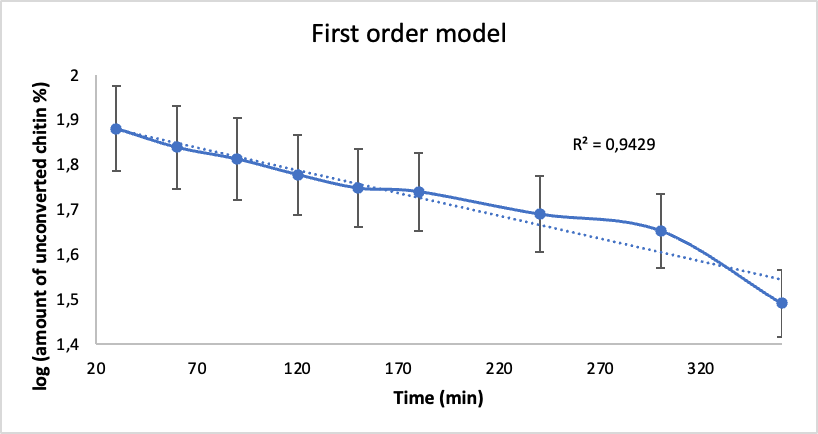

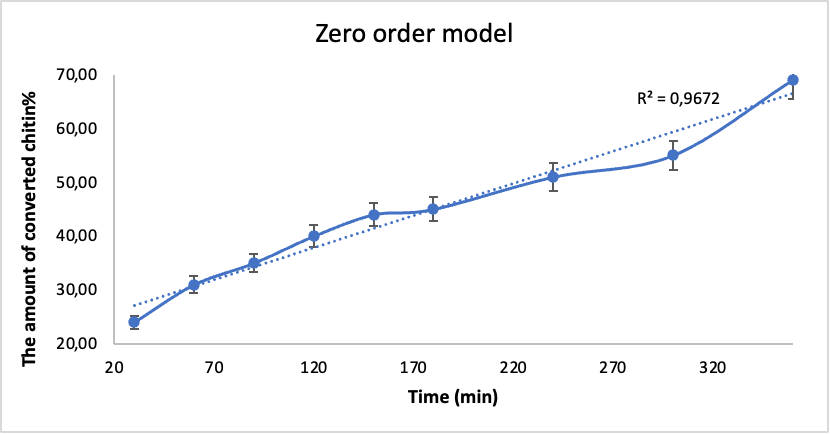

Supplementary Figure S1. Controlled release kinetic fits. Mathematical modeling of cumulative chitin hydrolysis by starch-coated silica nanoparticle formulations: (A) Korsmeyer-Peppas release profile model (R^2 = 0.9764); (B) Higuchi porous matrix diffusion plot (R^2 = 0.9636); (C) First-Order logarithmic decay fit (R^2 = 0.9429); (D) Zero-Order linear release model (R^2 = 0.9672). Error bars indicate standard deviations (n=3).
